# Fly stage trypanosomes recycle glucose catabolites and TCA cycle intermediates to stimulate growth in near physiological conditions

**DOI:** 10.1101/2020.12.17.423221

**Authors:** Oriana Villafraz, Marc Biran, Erika Pineda, Nicolas Plazolles, Edern Cahoreau, Rodolpho Ornitz Oliveira Souza, Magali Thonnus, Stefan Allmann, Emmanuel Tetaud, Loïc Rivière, Ariel M. Silber, Michael P. Barrett, Alena Zíková, Michael Boshart, Jean-Charles Portais, Frédéric Bringaud

## Abstract

*Trypanosoma brucei*, a protist responsible for human African trypanosomiasis (sleeping sickness), is transmitted by the tsetse fly, where the procyclic forms of the parasite develop in the proline-rich (1-2 mM) and glucose-depleted digestive tract. Proline is essential for the midgut colonization of the parasite in the insect vector, however other carbon sources could be available and used to feed its central metabolism. Here we show that procyclic trypanosomes can consume and metabolize metabolic intermediates, including those excreted from glucose catabolism (succinate, alanine and pyruvate), with the exception of acetate, which is the ultimate end-product excreted by the parasite. Among the tested metabolites, tricarboxylic acid (TCA) cycle intermediates (succinate, malate and α-ketoglutarate) stimulated growth of the parasite in the presence of 2 mM proline. The pathways used for their metabolism were mapped by proton-NMR metabolic profiling and phenotypic analyses of a dozen RNAi and/or null mutants affecting central carbon metabolism. We showed that (*i*) malate is converted to succinate by both the reducing and oxidative branches of the TCA cycle, which demonstrates that procyclic trypanosomes can use the full TCA cycle, (*ii*) the enormous rate of α-ketoglutarate consumption (15-times higher than glucose) is possible thanks to the balanced production and consumption of NADH at the substrate level and (*iii*) α-ketoglutarate is toxic for trypanosomes if not appropriately metabolized as observed for an α-ketoglutarate dehydrogenase null mutant. In addition, epimastigotes produced from procyclics upon overexpression of RBP6, showed a growth defect in the presence of 2 mM proline, which is rescued by α-ketoglutarate, suggesting that physiological amounts of proline are not sufficient *per se* for the development of trypanosomes in the fly. In conclusion, these data show that trypanosomes can metabolize multiple metabolites, in addition to proline, which allows them to confront challenging environments in the fly.

**Author Summary:** In the midgut of its insect vector, trypanosomes rely on proline to feed their energy metabolism. However, the availability of other potential carbon sources that can be used by the parasite is currently unknown. Here we show that tricarboxylic acid (TCA) cycle intermediates, *i.e.* succinate, malate and α-ketoglutarate, stimulate growth of procyclic trypanosomes incubated in medium containing 2 mM proline, which is in the range of the amounts measured in the midgut of the fly. Some of these additional carbon sources are needed for the development of epimastigotes, which differentiate from procyclics in the midgut of the fly, since their growth defect observed in the presence of 2 mM proline is rescued by addition of α-ketoglutarate. In addition, we have implemented new approaches to study a poorly explored branch of the TCA cycle converting malate to α-ketoglutarate, which was previously described as non-functional in the parasite, regardless of the glucose levels available. The discovery of this branch reveals that a full TCA cycle can operate in procyclic trypanosomes. Our data broaden the metabolic potential of trypanosomes and pave the way for a better understanding of the parasite’s metabolism in various organ systems of the tsetse fly, where it evolves.

## Introduction

*Trypanosoma brucei* is a hemoparasitic unicellular eukaryote sub-species of which cause Human African Trypanosomiasis (HAT), also known as sleeping sickness. The disease, fatal if untreated, is endemic in 36 countries in sub-Saharan Africa, with about 70 million people living at risk of infection [1]. The *T*. *brucei* life cycle is complex and the parasite must adapt to several dynamic environments encountered both in the insect vector, tsetse fly, and in the mammalian hosts. This is accomplished by cellular development, including adaptations of energy metabolism. Here, we focus on the insect stages of the parasite, the midgut procyclic (PCF) and epimastigote forms, by providing a comprehensive analysis of the carbon sources capable of feeding its central metabolism in tissue culture.

In glucose-rich mammalian blood, the metabolism of *T*. *brucei* bloodstream forms (BSF) relies on glucose, with most of the glycolytic pathway occurring in specialized peroxisomes called glycosomes [2]. BSF convert glucose, their primary carbon source, to the excreted end-product pyruvate, although low but significant amounts of acetate, succinate and alanine are also produced [3,4]. In contrast, the PCF mainly converts glucose to excreted acetate and succinate, in addition to smaller amounts of alanine, pyruvate and lactate [5]. Although PCF trypanosomes prefer to use glucose to support their central carbon metabolism *in vitro* [6,7], they rely on proline metabolism in the fly midgut [8]. This is due to the presumed low abundance, or absence, of free glucose in the hemolymph and tissues of the insect vector, which rely on amino acid for their own energy metabolism [9]. In this particular *in vivo* context, the PCF has developed an energy metabolism based on proline, which is converted to alanine, succinate and acetate (see Fig 1A) [7,10].

**Fig 1.**
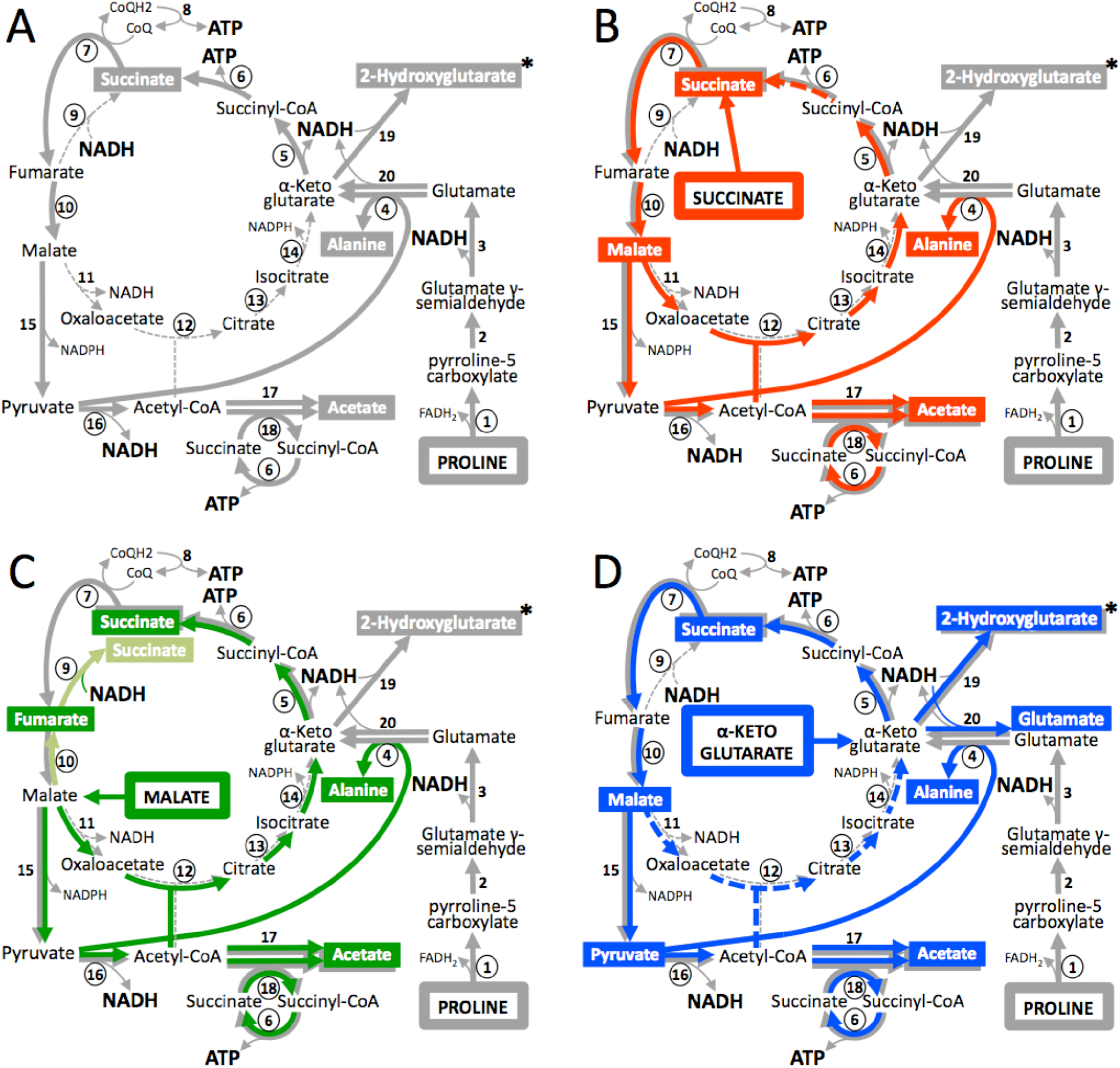
Proline metabolism of the PCF trypanosomes in the presence of other carbon sources. Panels A, B, C and D correspond to schematic metabolic representations of PCF trypanosomes incubated in glucose-depleted medium, containing proline (grey arrows) without or with succinate (red arrows), malate (green arrows) and α-ketoglutarate (blue arrows), respectively. End-products excreted from catabolism of proline and the other carbon sources are shown in rectangles with the corresponding colour code and the enzyme numbers under catabolism of succinate and α-ketoglutarate, respectively. In panel C, enzymatic reactions occurring in the reducing direction of the TCA cycle are shown in light green. The production and consumption of ATP and NAD^+^ are indicated in bold and the asterisks mean that production of 2-hydroxyglutarate from proline has not been previously described for PCF. It should be noted that, according to the literature, the TCA cycle does not work as a cycle in trypanosomes and only branches are used through anaplerotic reactions [16]. Indicated enzymes are : 1, proline only branches are used through anaplerotic reactions [16]. Indicated enzymes are : 1, proline (P5CDH); 4, alanine aminotransferase (AAT); 5, α-ketoglutarate dehydrogenase complex (KDH); 6, succinyl-CoA synthetase (SCoAS); 7, succinate dehydrogenase (SDH, complex II of the respiratory chain); 8, respiratory chain and mitochondrial ATP synthetase (oxidative phosphorylation); 9, mitochondrial NADH-dependent fumarate reductase (FRDm1); 10, mitochondrial fumarase (FHm); 11, mitochondrial malate dehydrogenase (MDHm); 12, citrate synthase (CS); 13, aconitase (ACO); 14, mitochondrial isocitrate dehydrogenase (IDHm); 15, mitochondrial malic enzyme (MEm); 16, pyruvate dehydrogenase complex (PDH); 17, acetyl-CoA thioesterase (ACH); 18, acetate:succinate CoA-transferase (ASCT); 19, unknown enzyme; 20, possibly NADH-dependent glutamate dehydrogenase.

As in other organisms, *T. brucei* PCF catabolism of proline is achieved by oxidation to glutamate, through proline dehydrogenase (PRODH, step 1 in Fig 1) [6] and pyrroline-5 carboxylate dehydrogenase (P5CDH, step 3) [8]. RNAi-mediated downregulation of *P5CDH* expression (^*RNAi*^P5CDH) is lethal for the PCF grown in glucose-depleted medium containing 6 mM proline (glucose-depleted conditions) and abolishes fly infections. This demonstrated that proline is essential for growth and development of insect-stage trypanosomes *in vivo* [8]. Proline derived glutamate is then converted, by alanine aminotransferase (AAT, step 4), to the TCA cycle intermediate α-ketoglutarate [11], which is further metabolized to succinate (steps 5-6) and malate (steps 7 and 10) *via* the TCA cycle enzymes working in the oxidative direction [7].

Alternatively, glutamate dehydrogenase, which catalyzes an oxidative deamination of glutamate, could also be involved in the production of α-ketoglutarate, as proposed for *T. cruzi*, but this has never been demonstrated in *T. brucei* [12]. Malate is then converted to pyruvate by the malic enzymes (step 15) [13], and serves as a substrate, together with glutamate, for AAT to produce α-ketoglutarate and secreted alanine (step 4). Alternatively, pyruvate is converted by the pyruvate dehydrogenase complex (PDH, step 16) to acetyl-CoA, which is further metabolized into the excreted end-product acetate by two redundant enzymes, *i.e.* acetyl-CoA thioesterase (ACH, step 17) and acetate:succinate CoA-transferase (ASCT, step 18) [14]. It is noteworthy that oxidation of acetyl-CoA through the TCA cycle (dashed lanes in Fig 1A), initiated by production of citrate by citrate synthase (CS, step 12), has not been detected so far in *T. brucei* [15]. Consequently, it is currently considered that the TCA cycle does not work as a cycle in trypanosomes, which only use branches of the TCA cycle fed by anaplerotic reactions [16].

The essential role of proline metabolism to trypanosomes in the insect vector has been well established [8]. However, the utilization of other carbon sources that may be available in the digestive tract and other organs of the insect, as well as the pathways of central carbon metabolism used by trypanosomes *in vivo*, are currently unknown or poorly understood. For instance, the transient availability of glucose for trypanosomes directly following tsetse blood meals could contribute significantly to parasite development in the midgut of the fly. In addition, possible metabolic interactions between trypanosomes and intestinal symbiotic bacteria might also be taken into account, as illustrated by the positive correlation between the presence of facultative symbionts including *Wrigglesworthia glossinidia* and *Sodalis glossinidius* and the ability of the tsetse fly to be infected by *T. brucei* [17,18]. Indeed these bacteria have been proposed to create a metabolic symbiosis with *T. brucei* through provision of precursors to threonine biosynthesis [19].

Here we show that PCF metabolizes carbon sources other than glucose and proline, such as succinate, alanine, pyruvate, malate and α-ketoglutarate. Interestingly, the TCA cycle intermediates succinate, malate and α-ketoglutarate stimulate growth of the parasites in *in vivo*-like conditions (2 mM proline [20], without glucose). We also took advantage of the high metabolic rate of these TCA cycle intermediates, to study the metabolic capacity of the parasite. This approach provided the first evidence for a complete canonical TCA cycle in PCF trypanosomes.

## Results

### Procyclic trypanosomes can re-metabolize end-products excreted from glucose degradation

To study the capacity of trypanosomes to metabolize unexpected carbon sources, such as those possibly excreted from the metabolism of the fly microbiota, we developed a model in which procyclic trypanosomes (PCF) have the possibility to re-consume partially oxidized metabolites excreted from their own glucose catabolism. It is noteworthy that this model may also represent an *in vivo* situation, when established PCF are exposed to a new blood meal of the insect. In this experimental model, the parasites were incubated at a high density in PBS containing low amounts of ^13^C-enriched glucose ([U-^13^C]-glucose, 0.5 mM) in the presence or absence of 4 mM proline. The production and possible re-consumption of excreted end-products from the metabolism of [U-^13^C]-glucose and/or non-enriched proline was monitored by analyzing the medium over-time during a 6 h incubation period using the ^1^H-NMR profiling approach. This quantitative ^1^H-NMR approach was previously developed to distinguish between [^13^C]-enriched and non-enriched excreted molecules produced from [^13^C]-enriched and non-enriched carbon sources, respectively [21–23]. In these conditions, all glucose is consumed within the first 1.5-2 h (Fig 2A-B). When [U-^13^C]-glucose is the only carbon source, PCF convert it to ^13^C-enriched acetate, succinate, alanine and pyruvate, in addition to lower amounts of non-enriched metabolites produced from an unknown internal carbon source (ICS) [21–23]. After 1 h of incubation, net production of ^13^C-enriched succinate and pyruvate stopped while glucose remained in the medium. Interestingly, ^13^C-enriched acetate was still excreted even after glucose depletion, suggesting that acetate was produced from the metabolism of excreted end-products, such as succinate and pyruvate (Fig 2A, top). The same pattern was observed from the non-enriched excreted metabolites produced from the ICS (Fig 2A, low).

**Fig 2.**
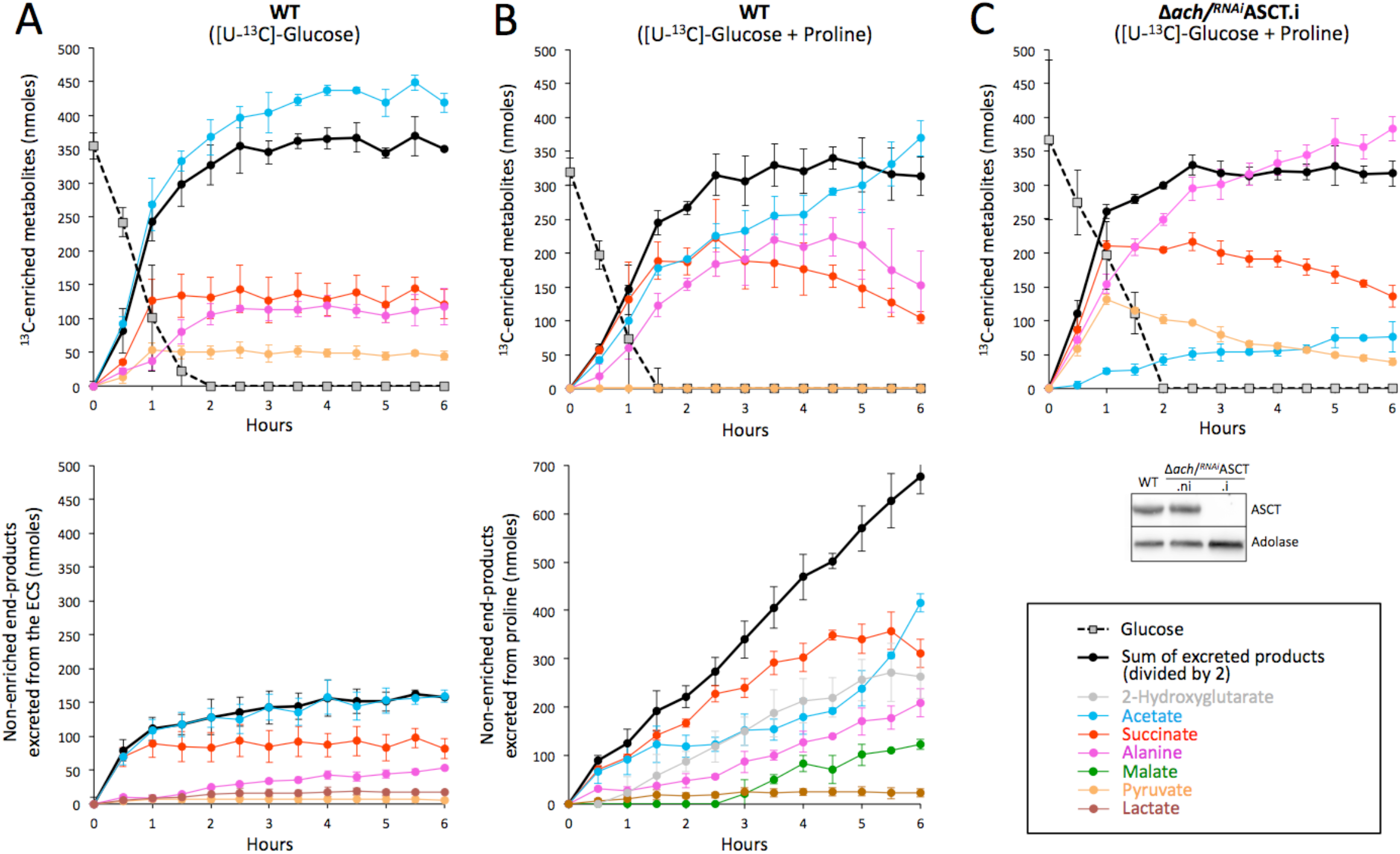
Kinetic analyses of end-products excretion from [U-^13^C]-glucose and proline metabolism. PCF cells were incubated for 6 h in PBS containing 0.5 mM [U-^13^C]-glucose in the presence (panel B) or not (panel A) of 4 mM non-enriched proline (Pro). Excreted end-products were analyzed in the spent medium every 0.5 h by ^1^H-NMR spectrometry. The top panels A and B show the amounts of ^13^C-enriched end-products excreted from [U-^13^C]-glucose metabolism and the amounts of glucose present in the spent medium. The bottom panels show the amounts of non-enriched end-products produced from the metabolism of the unknown internal carbon source (panel A) or from the metabolism of proline plus the unknown internal carbon source (panel B). In panel C, the kinetics of end-products excreted from 0.5 mM [U-^13^C]-glucose in the presence of 4 mM proline was determined for the Δ*ach*/^*RNAi*^ASCT.i mutant cell line as performed in panel B. Western blot controls with the anti-ASCT and anti-aldolase immune sera are shown below the graph.

Addition of proline strongly stimulated this re-utilization of glucose-derived succinate (Fig 2B, top). In these incubation conditions, glucose-derived pyruvate was no longer excreted since it is used as a substrate by alanine aminotransferase (AAT, step 4 in Fig 1A) to convert proline-derived glutamate to α-ketoglutarate [7]. Alanine was also re-used, although with a 2.5 h delay compared to succinate. This re-utilization was also seen for proline-derived succinate (Fig 2B, low) as proline-derived acetate increased at the expense of succinate after 4.5 h of incubation. Combined, these data suggested that glucose-derived and/or proline-derived succinate, pyruvate and to a lower extent alanine can be re-utilized and converted to acetate.

Since acetate is produced by both the acetate:succinate CoA-transferase (ASCT, step 18 in Fig 1A) and the acetyl-CoA thioesterase (ACH, step17), the role of the acetate branch in the further metabolism of excreted end-products was addressed. The same time-course was conducted on a Δ*ach*/^*RNAi*^ASCT double mutant, in which ASCT expression was knocked down by RNAi in the null ACH background. After 2 days of incubation, the tetracycline induced Δ*ach*/^*RNAi*^ASCT (Δ*ach*/^*RNAi*^ASCT.i) cell line showed an 80% reduction in acetate production from glucose metabolism, compared to the parental cell line (Fig 2B-C), which is consistent with the 90% reduction in acetate excretion previously described for this cell line [21]. The residual acetate production was attributed to incomplete downregulation of ASCT expression, although ASCT could not be detected by Western blot [21] (Fig 2C). The excretion of glucose-derived pyruvate in the Δ*ach*/^*RNAi*^ASCT.i mutant relates to the limited capacity of the acetate branch. Interestingly, succinate and pyruvate excreted during the first hour of incubation are re-consumed and converted to acetate and alanine, indicating that alanine is the ultimate excreted end-product when the acetate branch is limiting (Fig 2C).

### Succinate, pyruvate and alanine are metabolized in the presence of glucose or proline

To determine how the main end-products excreted from glucose catabolism (succinate, alanine, pyruvate or acetate) are metabolized in the presence or absence of glucose or proline, we analyzed, by quantitative ^1^H-NMR, the exometabolome of PCF incubated with [U-^13^C]-succinate, [U-^13^C]-alanine, [U-^13^C]-pyruvate or [U-^13^C]-acetate in the presence or absence of equal amounts of non-enriched glucose or proline. Succinate was poorly consumed alone, however, the presence of glucose or proline stimulates its consumption by 3.6- and 4.6-fold, respectively (Fig 3A). This is consistent with the increased conversion of glucose-derived succinate to acetate, in the presence of proline (Fig 2A-B). The quantities of excreted end-products from proline catabolism were reduced only 8% in the presence of succinate, compared to a 53% reduction in the presence of glucose (Fig 3A). This suggests that, in contrast to the previously reported glucose-induced downregulation of proline metabolism [6,7], succinate addition does not limit the rate of proline metabolism. In the presence of [U-^13^C]-proline, succinate is converted to malate, acetate, alanine and traces of fumarate, which represent 40.5%, 43.2%, 16.4% and 1.5% of the excreted end-products, respectively (Fig 3B). According to the current metabolic model, succinate enters the tricarboxylic acid (TCA) cycle where it is converted to fumarate by succinate dehydrogenase (SDH, step 7 in Fig 1B) and to malate by the mitochondrial fumarase (FHm, step 10). The malic enzymes then produce pyruvate (MEm, step 15 and the cytosolic MEc not shown), which feeds the AAT (step 4) for production of alanine, or the pyruvate dehydrogenase (PDH) complex plus the ACH/ASCT steps (steps 17 and 18) for production of acetate. In agreement with this model, extracellular succinate and proline-derived succinate were no longer metabolized to acetate in the ^*RNAi*^SDH.i cell line (Fig 3B). As expected, acetate production from succinate, as well as from proline, was abolished in the tetracycline-induced PDH subunit E2 RNAi mutant cell line (^*RNAi*^PDH-E2.i). The effective block of PDH activity is reflected by the increased excretion of succinate-derived pyruvate, the substrate of the PDH complex (Fig 3B).

**Fig 3.**
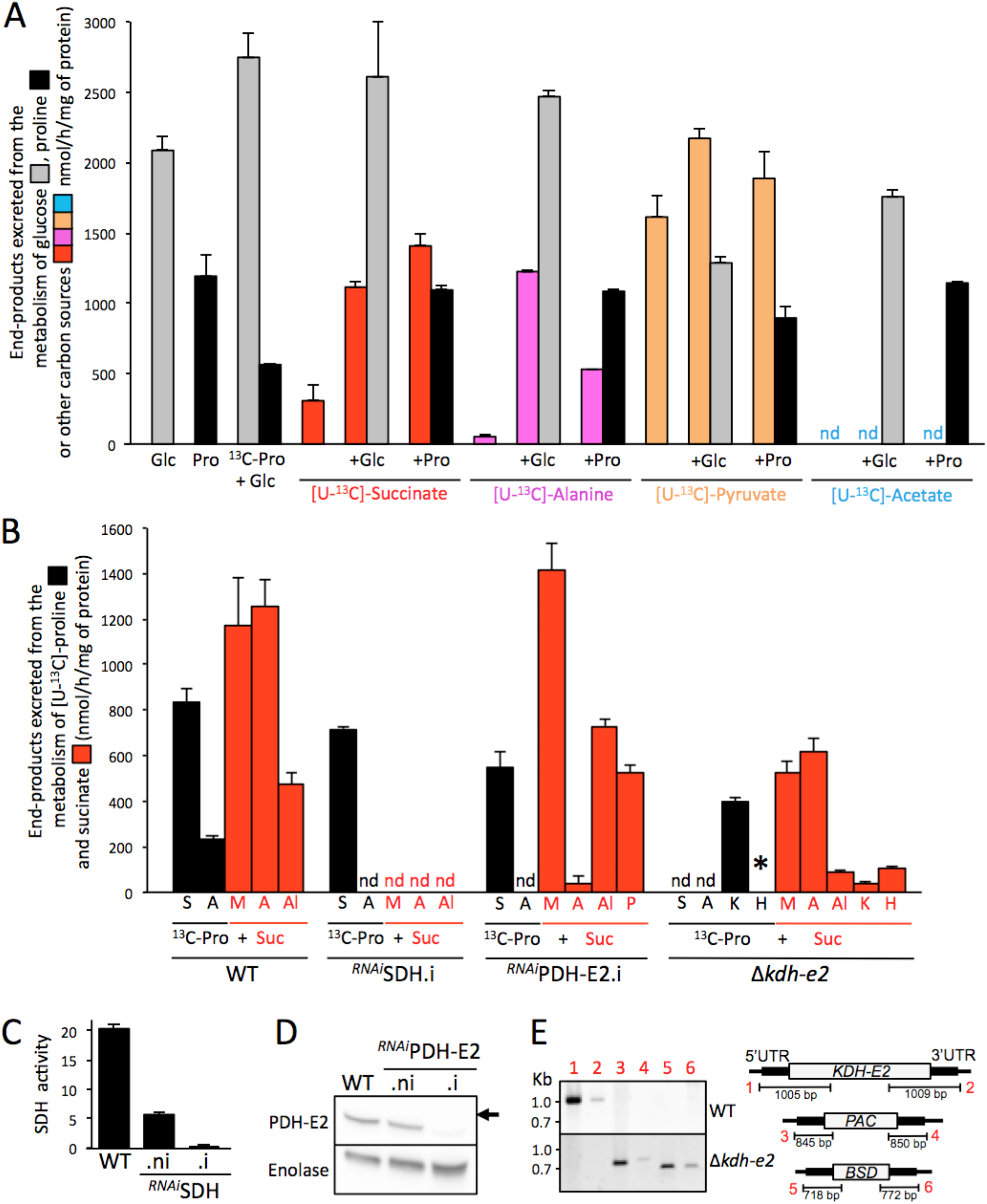
Proton (^1^H) NMR analyses of end-products excreted from the metabolism of ^13^C-enriched succinate, alanine, pyruvate and acetate. In panel A, PCF trypanosomes were incubated for 6 h in PBS containing 4 mM [U-^13^C]-succinate, [U-^13^C]-alanine, [U-^13^C]-pyruvate or [U-^13^C]-acetate alone or in combination with 4 mM glucose (+Glc) or proline (+Pro) before analysis of the spent medium by ^1^H-NMR spectrometry. As reference, the same experiment was performed with 4 mM proline (Pro), 4 mM glucose (Glc) or 4 mM glucose with 4 mM [U-^13^C]-proline (^13^C-Pro). The amounts of each end-product excreted are documented in S1 Table. Panel B shows equivalent ^1^H-NMR spectrometry experiments performed on the parental (WT), ^*RNAi*^SDH.i, ^*RNAi*^PDH-E2.i and Δ*kdh-e2* cell lines incubated with 4 mM [U-^13^C]-proline and 4 mM succinate. Because of high background, ^13^C-enriched 2-hydroxyglutarate produced from [U-^13^C]- proline cannot be quantified, however, it is detectable in the Δ*kdh-e2* cell line as indicated by an asterisk (*). Traces of fumarate produced from these carbon sources are not shown in the figure. Abbreviations: A, acetate; Al, alanine; H, 2-hydroxyglutarate; K, α-ketoglutarate; M, malate; P, pyruvate; S, succinate; nd, not detectable; *, detectable but not quantifiable. The efficiency of RNAi-mediated downregulation of SDH and PDH-E2 expression in the tetracycline-induced (.i) or non induced (.ni) cell line, as well as in the parental cell line (WT), was determined by SDH activity assays (panel C) and Western blotting with the anti-PDH-E2 and anti-enolase (control) immune sera (panel D). Panel E shows a PCR analysis of genomic DNA isolated from the parental (WT) and Δ*kdh-e2* cell line. Lanes 1 to 6 of the gel picture correspond to different PCR products described in the right panel. As expected, PCR amplification of the *KDH-E2* gene (lanes 1-2) was only observed in the parental cell line, while *PAC* and *BSD* PCR-products were observed only in the Δ*kdh-e2* mutant (lanes 3-4 and 5-6, respectively).

[U-^13^C]-Alanine was poorly metabolized alone, but addition of glucose or proline considerably stimulated its consumption, with the production of ^13^C-enriched end-products being 23-fold and 10-fold increased, respectively (Fig 3A). ^13^C-enriched acetate represents 100% and 82% of the excreted end-products from [U-^13^C]-Alanine in the presence of proline and glucose, respectively (S1 Table). [U-^13^C]-Pyruvate was converted to ^13^C-enriched alanine and acetate in the presence or absence of glucose or proline. The rate of end-product excretion was only increased by 35% and 17% in the presence of glucose and proline, respectively (Fig 3A). In contrast, no ^13^C-enriched molecules were detected by ^1^H-NMR in the exometabolome of PCF incubated with [U-^13^C]-acetate, in the presence or absence of glucose or proline, suggesting that acetate is not further metabolized through the central metabolism (Fig 3A). Together, these data show that acetate is the ultimate excreted end-product of the metabolism of glucose and other carbon sources, while succinate, pyruvate and alanine, which are also excreted, can be re-utilized and converted to acetate.

### TCA cycle intermediates stimulate growth of the PCF in *in vivo*-like conditions

The midgut of the tsetse fly, the natural environment of PCF, lacks glucose between blood meals and contains proline in the low mM range (1-2 mM) [20], which is used by the parasite for its energy metabolism [8]. To determine if succinate, alanine, acetate, pyruvate and lactate stimulate growth of the parasite in insect-like conditions, we estimated the growth of the PCF trypanosomes as a function of increasing concentrations of these metabolites in the glucose-depleted SDM79 medium containing 2 mM proline, using the Alamar Blue assay, as previously described [23]. Among them, succinate (1 to 10 mM) was able to stimulate growth, with a maximum effect at 10 mM, while pyruvate showed a moderate effect (S1 Fig). This succinate-dependent growth stimulation is observed in the presence of up to 2 mM proline, but not in high-proline conditions (12 mM) (Fig 4A). As controls, 1 to 20 mM proline stimulated growth in the presence 0.2 mM and 2 mM proline, but not in the presence of 12 mM proline. Among six other TCA cycle intermediates tested plus glutamate and aspartate, malate and α-ketoglutarate also stimulated growth in the presence of 2 mM proline (and 0.2 mM), with a maximum effect on growth also at 10 mM (S1 Fig and Fig 4A). This growth stimulation was confirmed by performing growth curves on cells maintained in the exponential growth phase (between 10^6^ and 10^7^ cells/ml) over 15 days in the presence of 2 mM proline and 10 mM of succinate, malate or α-ketoglutarate (Fig 4B). In the presence of these metabolites the culture doubling time was reduced by approximately 1.2 fold, with an increased growth rate compared to that using 2 mM proline alone. As observed for succinate, NMR analyses of excreted end-products from the metabolism of malate and α-ketoglutarate showed that the addition of equimolar amounts of [U-^13^C]-proline induced an increase of the rate of malate and α-ketoglutarate consumption by 5.7 and 6.6 fold, respectively (Fig 4C). 6.9 and 2.8 fold increases of malate and α-ketoglutarate consumption were seen with [U-^13^C]-glucose, respectively.

**Fig 4.**
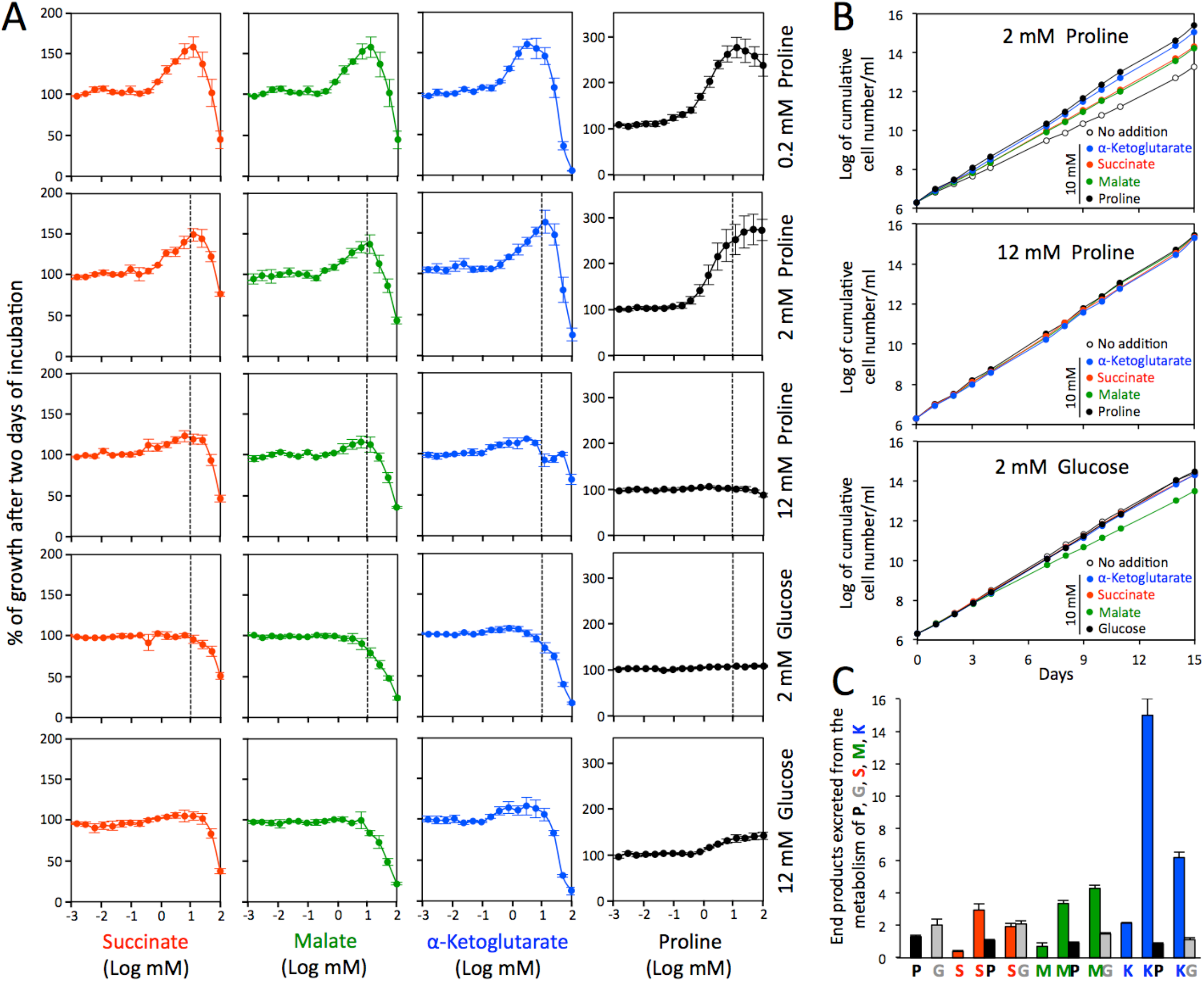
Succinate, malate and α-ketoglutarate stimulate growth of the PCF. Panel A shows growth of the PCF trypanosomes in the glucose-depleted SDM79 medium containing 0.2 mM, 2 mM (low-proline) or 12 mM (high-proline) proline, 2 mM glucose/2 mM proline (low-glucose) or 12 mM glucose/2 mM proline (high-glucose) in the presence of added 10 μM to 100 mM succinate, malate, α-ketoglutarate or proline, using the Alamar Blue assay. Incubation was started at 2 × 10^6^ cell density and the Alamar Blue assay was performed after 48 h at 27°C. The dashed line indicates the concentrations of succinate, malate, α-ketoglutarate or proline (10 mM) used in panel B, which shows growth curves of the PCF in low-proline, high-proline and low-glucose conditions in the presence or not of 10 mM of each metabolite. Cells were maintained in the exponential growth phase (between 10^6^ and 10^7^ cells/ml), and cumulative cell numbers reflect normalization for dilution during cultivation. In panel C, the PCF trypanosomes were incubated for 6 h in PBS containing 4 mM succinate (S), malate (M) or α-ketoglutarate (K), in the presence or absence of 4 mM [U-^13^C]-proline (P) or [U-^13^C]-glucose (G), before analysis of the spent medium by ^1^H-NMR spectrometry. As a control, the cells were also incubated with 4 mM [U-^13^C]-proline (P) or [U-^13^C]-glucose (G) alone. The amounts of end-products excreted from the metabolism of proline (black), glucose (grey), succinate (red), malate (green) and α-ketoglutarate (blue) are expressed as μmol excreted/h/mg of protein.

The effect of succinate, malate and α-ketoglutarate was also determined on PCF grown in glucose-rich conditions. At most, succinate, α-ketoglutarate and the proline control have a minor stimulatory effect in the presence of 2 mM or 12 mM glucose (2 mM proline) (Figs 4A-B). However, addition of 10 mM malate to PCF grown in the presence 2 mM glucose slightly slowed growth of the parasite (Fig 4B). Malate was consumed 22% more in glucose-rich than in glucose-depleted conditions and its presence induced a 27% reduction of glucose consumption (Fig 4C), suggesting that switch to partial malate metabolism is less efficient than catabolism of glucose alone.

### The TCA cycle is used to metabolize malate in procyclic trypanosomes

^1^H-NMR spectrometry analyses showed that, in the presence of proline, malate is converted in almost equal amounts to fumarate and succinate (35.9% and 38.6% of the excreted end-products), in addition to alanine and acetate (14.9% and 10.6% of the excreted end-products) (Fig 5A). According to the current view, malate is converted by the malic enzymes to pyruvate (step 15 in Fig 1C), a precursor for the production of alanine and acetate, as described above (steps 4 and 16-18). As observed for the metabolism of succinate, production of acetate from malate is abolished in the ^*RNAi*^PDH-E2.i cell line (Fig 5A), with an accumulation of malate-derived pyruvate and alanine. Malate can also be converted by FHm to fumarate (step 10), which is further reduced to succinate by the mitochondrial NADH-dependent fumarate reductase (FRDm1, step 9). It is to note that the cytosolic (FHc) and glycosomal (FRDg) isoforms of these two enzymes, respectively, could also be involved in succinate production [24,25], justifying the use of the ^*RNAi*^FRDg/m1.i and ^*RNAi*^FHc/m.i double mutants to study the production of succinate from malate. Succinate production from malate was reduced but not abolished in either of these two double mutants, suggesting that PCF uses an alternative pathway in addition to this reducing branch (Fig 5A). Indeed, malate is also reduced to succinate by TCA cycle enzymes (steps 11-14 and 5-6), as inferred by diminished secretion of malate-derived succinate in the Δ*aco* (aconitase, step 13) and Δ*kdh-e2* (α-ketoglutarate dehydrogenase subunit E2, step 5) mutant cell lines (Fig 5A). Two other lines of evidence supported the utilization of the oxidative branch of the TCA cycle to produce succinate. First, the Δ*kdh-e2* null mutant excreted α-ketoglutarate from malate metabolism. Second, the expected abolition of fumarate production from malate in the ^*RNAi*^FHc/m.i double mutant (Fig 5A), implies that the malate-derived succinate cannot be produced by the fumarate reductase activity, but by the TCA cycle activity. The relatively high flux of malate consumption is probably the consequence of an efficient maintenance of the mitochondrial redox balance, with NADH molecules produced in the oxidative branches (succinate production through the TCA cycle and acetate production) being reoxidized, at least in part, by the reductive branch (succinate production by fumarate reductases). This resembles the malate dismutation phenomenon well described in anaerobic parasites (for reviews see [26,27]). It is of note that, in the presence of proline, succinate was also converted to succinyl-CoA and probably to succinate through the TCA cycle as inferred by the accumulation of excreted non-enriched α-ketoglutarate in the Δ*kdh-e2* null mutant incubated with succinate and [U-^13^C]-proline (Figs 3B and 1B). These data confirmed that the TCA cycle operates as a complete cycle in these growth conditions.

**Fig. 5.**
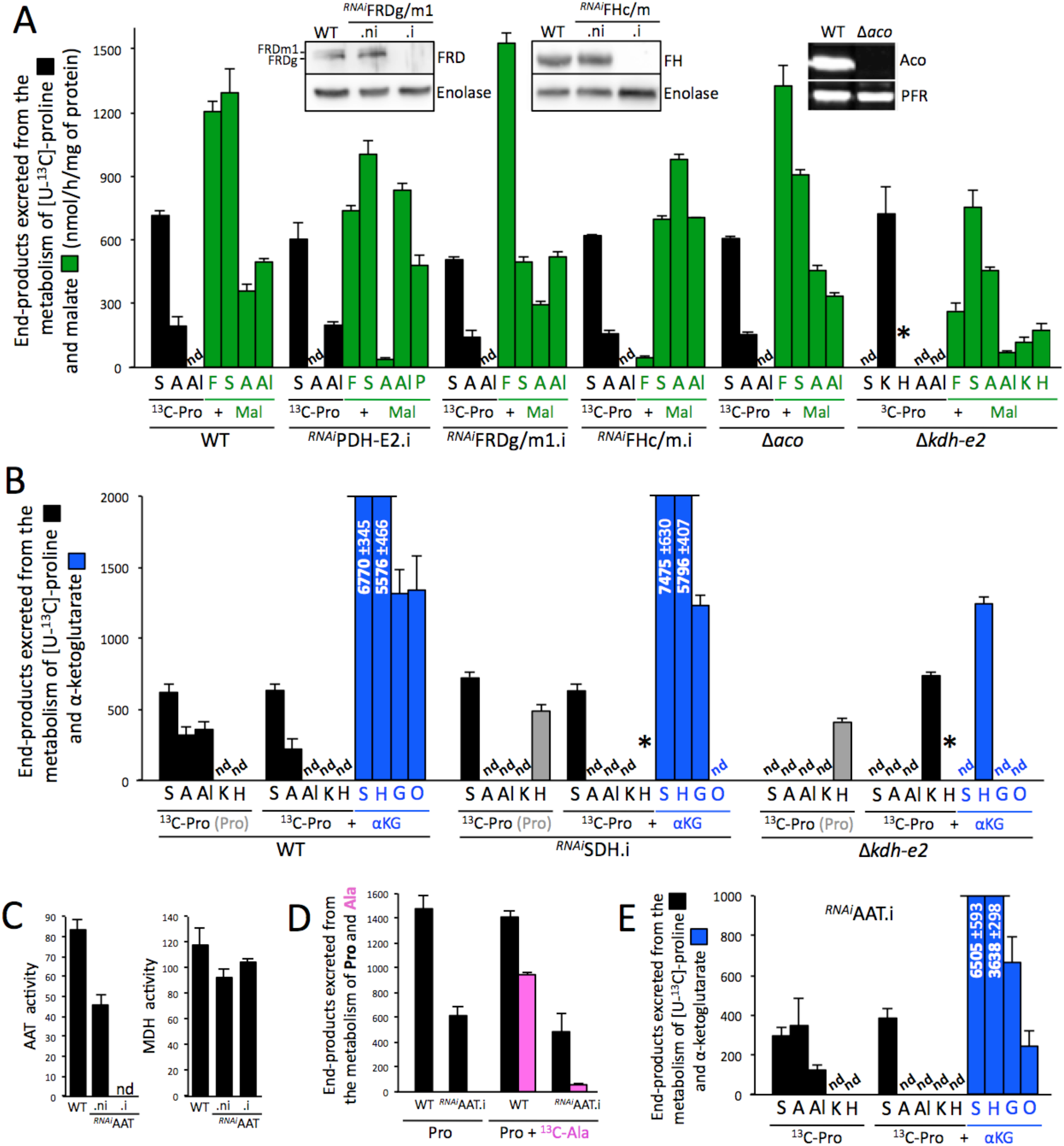
^1^H-NMR analyses of end-products excreted from the metabolism of malate and α-ketoglutarate. Panel A shows ^1^H-NMR spectrometry analyses of the exometabolome produced by the parental (WT), ^*RNAi*^PDH-E2.i, ^*RNAi*^FRDg/m1.i, ^*RNAi*^FHc/m.i, Δ*aco* and Δ*kdh-e2* cell lines incubated with 4 mM [U-^13^C]-proline (^13^C-Pro) and 4 mM malate (Mal). The insets show Western blot analyses with the immune sera indicated in the right margin of the parental (WT), the tetracycline-induced (.i) and non-induced (.ni) ^*RNAi*^FRDg/m1 and ^*RNAi*^FHc/m cell lines, and the Δ*aco* cell line. Panel B is equivalent to panel A with the parental (WT), ^*RNAi*^SDH.i and Δ*kdh-e2* cell lines incubated in the presence of [U-^13^C]-proline alone (^13^C-Pro) or with α-ketoglutarate (^13^C-Pro + αKG). Because of high background, ^13^C-enriched 2-hydroxyglutarate produced from [U-^13^C]-proline cannot be quantified, however, asterisks (*) mean that it is detectable. Since the amounts of 2-hydroxyglutarate produced from non-enriched proline are quantifiable, the values are indicated with grey columns when applicable. The excreted amounts are indicated in the truncated columns (nmol/h/mg of protein). The AAT and MDH (control) enzymatic activities of the parental (WT) and the tetracycline-induced (.i) and non-induced (.ni) ^*RNAi*^AAT cell lines are shown in panel C. In panel D, the WT and ^*RNAi*^AAT.i cells were incubated for 6 h in PBS containing 4 mM proline (Pro) with or without 4 mM [U-^13^C]-alanine (^13^C-Ala), before analysis of the spent medium by ^1^H-NMR spectrometry. The amounts of end-products excreted from the metabolism of proline (black) and alanine (pink) are expressed as μmol excreted/h/mg of protein. Panel E is equivalent to panel B, except that the ^*RNAi*^AAT.i is analyzed. Abbreviations: A, acetate; Al, alanine; F, fumarate; G, glutamate; H, 2-hydroxyglutarate; K, α-ketoglutarate; nd, not detectable; M, Malate; O, malate + pyruvate + acetate + alanine; S, succinate; *, detected but not quantifiable.

### Metabolism of α-ketoglutarate in the presence of proline

To our surprise, ^1^H-NMR spectrometry analyses of α-ketoglutarate metabolism showed that its rate of consumption in the presence of equal amounts (4 mM) of proline (~15 μmole/h/mg of protein) is ~15-times higher compared to that of glucose (~1 μmole/h/mg of protein), the latter having previously been considered as the most rapidly degraded carbon source by PCF trypanosomes [6]. As observed for the metabolism of succinate and malate, the rate of α-ketoglutarate catabolism is increased in the presence of proline or glucose (Fig 4C). In the presence of proline, α-ketoglutarate is mainly converted to equivalent amounts of succinate and 2-hydroxyglutarate (45.5% and 37.2% of the excreted end-products, respectively), with significant amounts of glutamate (8.7%), acetate (3.6%), pyruvate (3.1%) and malate (1.6%), as well as less than 1% of alanine and lactate (Fig 5B). The production of 2-hydroxyglutarate and glutamate from α-ketoglutarate was validated by comparing the exometabolome of the PCF incubated with [U-^13^C]-proline or [U-^13^C]-proline/α-ketoglutarate with 2-hydroxyglutarate and glutamate standards (S2 Fig). According to the current model, succinate is produced from α-ketoglutarate by the successive action of α-ketoglutarate dehydrogenase (KDH, step 5 in Fig 1D) and succinyl-CoA synthetase (SCoAS, step 6). Succinate then feeds production of malate, pyruvate, alanine and acetate (Fig 1D, O for other), as evidenced by their absence in the exometabolome of the ^*RNAi*^SDH.i cell line incubated with [U-^13^C]-proline and α-ketoglutarate (Fig 5B).

The Δ*kdh-e2* null mutant excretes only 2-hydroxyglutarate from metabolism of α-ketoglutarate, suggesting that a single reduction step, catalyzed by an as yet unknown enzyme, produces 2-hydroxyglutarate from α-ketoglutarate [28,29]. It has recently been proposed that 2-hydroxyglutarate detected in the *T. brucei* BSF metabolome results from the promiscuous action of the NADH-dependent malate dehydrogenase on α-ketoglutarate [30]. 2-hydroxyglutarate is also produced from malate and succinate by the Δ*kdh-e2* null mutant (Figs 3B and 5A). In addition, revisiting NMR spectrometry data showed that 2-hydroxyglutarate is also excreted by the parental PCF from proline metabolism in the presence of glucose, but not in its absence (Fig 2B). It is noteworthy that ^13^C-enriched 2-hydroxyglutarate molecules produced from [U-^13^C]-proline are barely detectable due to high background and observed but not quantifiable in the ^*RNAi*^SDH.i and Δ*kdh-e2* cell lines (Figs 3B and 5).

Since, α-ketoglutarate is produced from glutamate by the AAT transamination reaction (step 4) [11], the ^*RNAi*^AAT.i cell line was studied to determine if AAT catalyzes the reverse reaction. The AAT activity was no longer detectable in the ^*RNAi*^AAT.i cell line (Fig 5C), however, the catabolism of proline was only 2.5-fold reduced in the ^*RNAi*^AAT.i cell line compared to the parental cell line (Fig 5D). In addition, the ^*RNAi*^AAT.i cell line still produced glutamate from α-ketoglutarate (Fig 5E). This suggests that an alternative enzyme is involved in the reversible conversion of glutamate to α-ketoglutarate. Interestingly, alanine conversion to acetate, which theoretically requires the AAT activity working in the direction of glutamate production (see Fig 1A), was 20-fold reduced in the ^*RNAi*^AAT.i cell line (Fig 5D), suggesting that the alternative enzyme is not using alanine as substrate to converts α-ketoglutarate to glutamate and is probably not another aminotransferase showing substrate promiscuity. Thus, the best candidate is the NADH-dependent glutamate dehydrogenase previously described in trypanosomatids [12,31].

It is noteworthy that the possible NADH consumption through reduction of α-ketoglutarate to 2-hydroxyglutarate (5.6 ±0.47 μmol produced/h/mg of protein) and glutamate (1.3 ±0.17 μmol produced/h/mg of protein) may compensate NADH production by the KDH reaction (6.8 ±0.35 μmol of succinate produced/h/mg of protein).

### α-Ketoglutarate rescued the growth defect of the ^*RNAi*^PRODH.i and ^*RNAi*^AAT.i mutants

We previously showed that proline dehydrogenase (PRODH), which catalyzes the first step of proline catabolism, is important for the growth of the PCF in glucose-depleted conditions [6]. Growth of the ^*RNAi*^PRODH.i cell line was considerably reduced but not abolished in glucose-depleted conditions, probably because of the residual PRODH activity (Fig 6A) and residual conversion of proline to excreted end-products (Fig 6B). As expected, proline did not rescue growth of the ^*RNAi*^PRODH.i mutant in glucose-depleted conditions. However, succinate and malate improved growth of the mutant and more importantly α-ketoglutarate completely rescued its growth (Fig 6C), even in the presence of 12 mM proline (Fig 6D). Unfortunately, we failed to obtain the Δ*prodh* null mutant in standard glucose-rich conditions, with or without 10 mM α-ketoglutarate, probably because minimal proline catabolism is required even in the presence of glucose. As mentioned above, proline metabolism was 2.5-fold reduced in the ^*RNAi*^AAT.i mutant cell line (Fig 5D). This caused a slight growth defect in glucose-depleted conditions that was rescued by the addition of α-ketoglutarate, as observed for the ^*RNAi*^PRODH.i mutant (Fig 7B). Collectively, these data suggest that metabolism of α-ketoglutarate, and to a lesser extent in the case of the ^*RNAi*^PRODH.i cell line, succinate and malate, compensate for the lack of proline metabolism. To confirm these data, the ability of these carbon sources to replace proline was tested under long-term growth conditions. Because of the auxotrophy of *T. brucei* for proline, which is necessary for protein biosynthesis, the growth medium contained 0.2 mM proline. In these conditions, the growth defect observed in the absence of an additional carbon source is partially rescued with the same efficiency by 10 mM α-ketoglutarate, glucose, succinate and malate (S3 Fig). Interestingly, the growth rate in low-proline conditions is the same whether in the presence of glucose or malate/succinate/α-ketoglutarate, thus confirming that these TCA cycle intermediates are excellent carbon and energy sources for PCF.

**Fig. 6.**
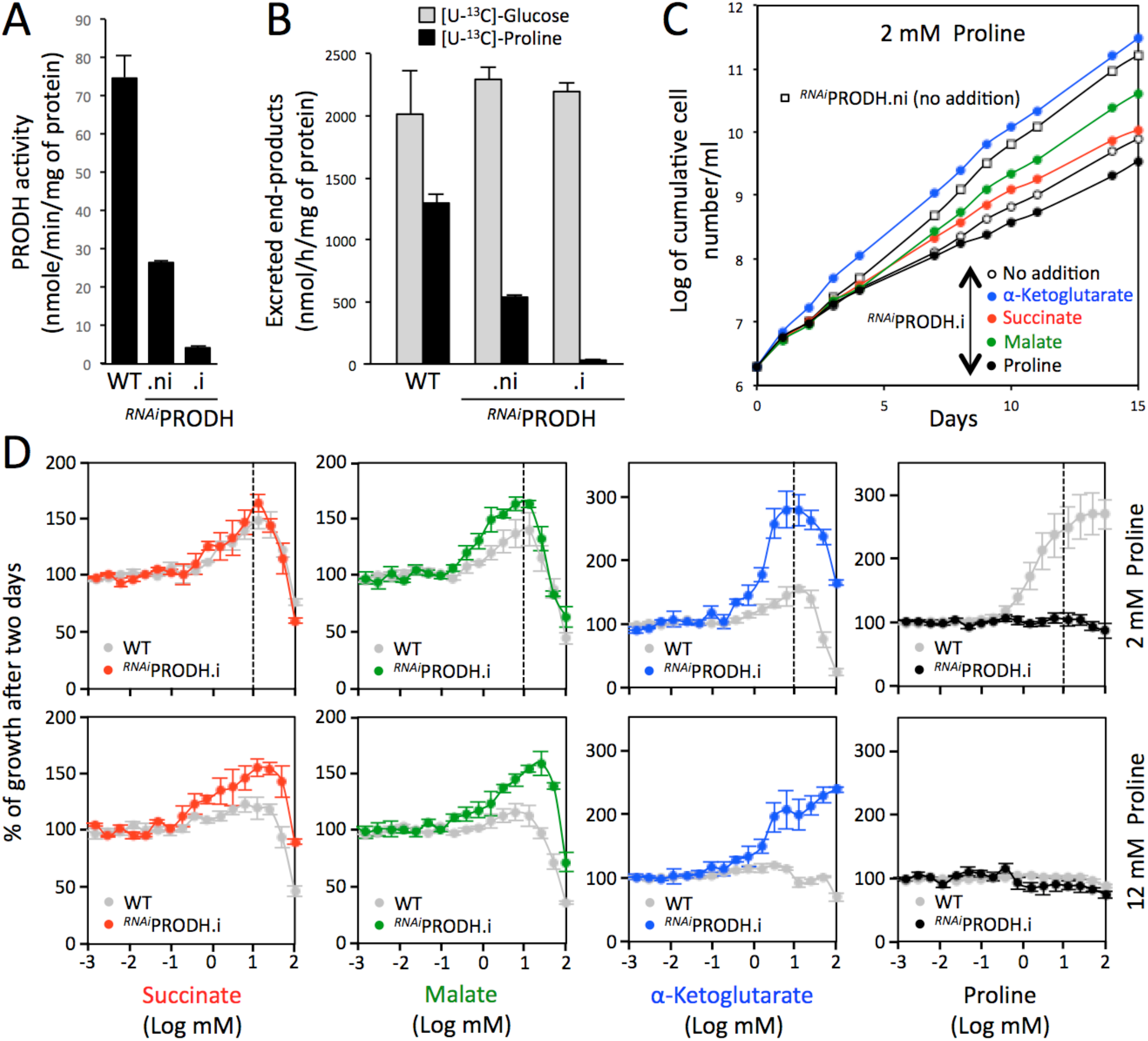
Growth of the ^*RNAi*^PRODH.i mutant is rescued by α-ketoglutarate. The efficiency of RNAi-mediated downregulation of PRODH expression in the tetracycline-induced (.i) or non induced (.ni) ^*RNAi*^PRODH cell line, as well as in the parental cell line (WT) was determined by the PRODH activity assay (panel A) and ^1^H-NMR quantification of end-products excreted from metabolism of proline and glucose in two independent experiments (panel B). Panel C shows growth curves of the ^*RNAi*^PRODH.i cell line in glucose-depleted medium containing 2 mM proline, in the presence (colored circles) or absence (open circles) of 10 mM α-ketoglutarate, malate, succinate or proline. Cells were maintained in the exponential growth phase and cumulative cell numbers reflect normalization for dilution during cultivation. The effect of 10 mM succinate, malate, α-ketoglutarate or proline on growth of the parental and ^*RNAi*^PRODH.i cell lines in glucose-depleted medium containing 2 mM proline, using the Alamar Blue assay described in Figure 4, is shown in panel D.

**Fig. 7.**
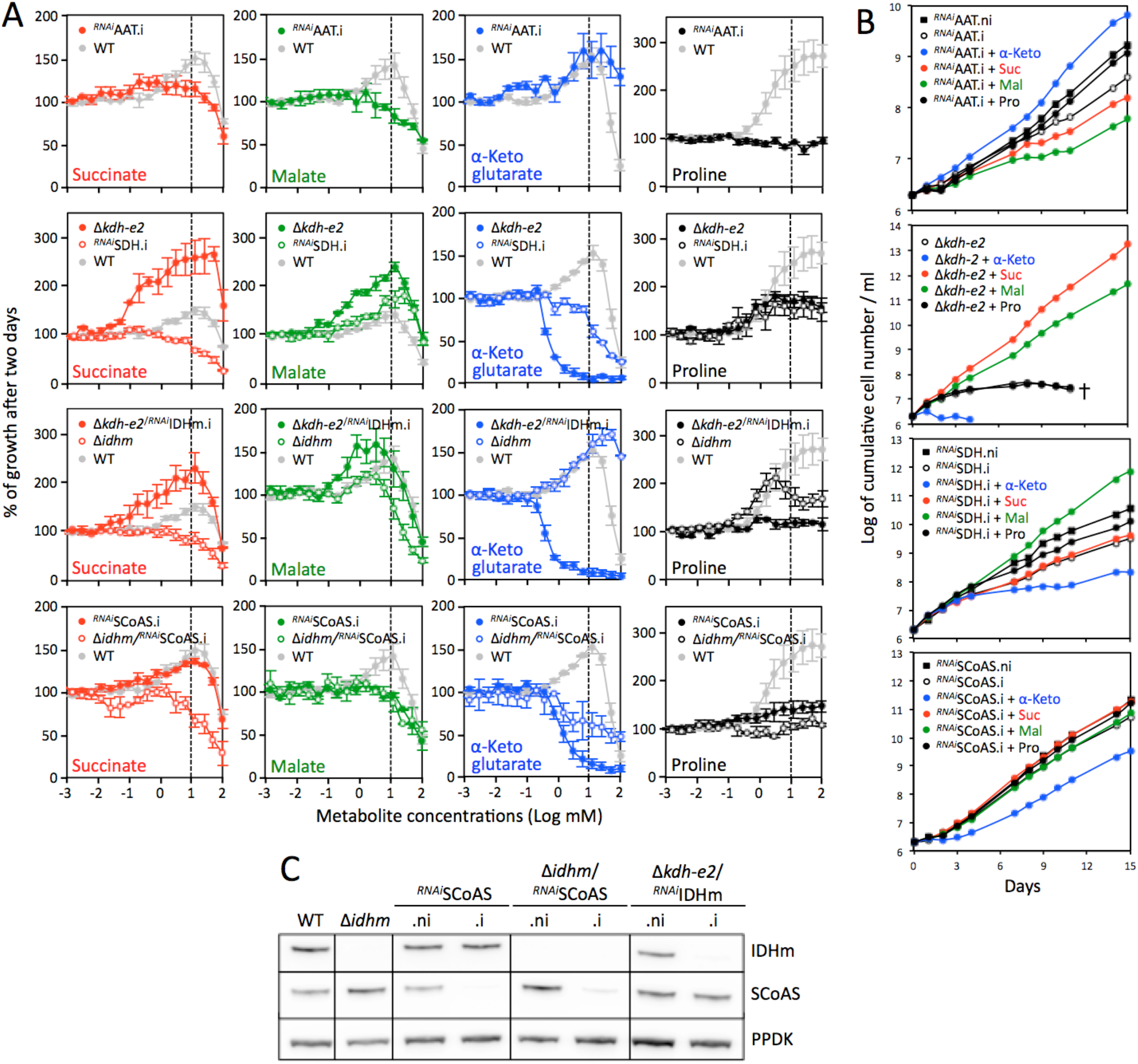
α-ketoglutarate is toxic for the Δ *kdh-e2*, ^*RNAi*^SCoAS.i and ^*RNAi*^SDH.i cell lines. Panel A compares the effect of succinate, malate, α-ketoglutarate or proline (10 mM) on growth of the parental and mutant cell lines in glucose-depleted medium containing 2 mM proline, using the Alamar Blue assay described in Figure 4. Panel B shows growth curves of the ^*RNAi*^AAT, Δ*kdh-e2*, ^*RNAi*^SDH and ^*RNAi*^SCoAS cell lines incubated in glucose-depleted medium containing 2 mM proline, in the presence (colored circles) or absence (open circles) of α-ketoglutarate, malate, succinate or proline (10 mM) (.i, tetracycline-induced cells; .ni, non induced cells). Cells were maintained in the exponential growth phase and cumulative cell numbers reflect normalization for dilution during cultivation. Panel C shows Western blot analyses with the immune sera indicated in the right margin of the parental (WT), Δ*idhm* and tetracycline-induced (.i) and non-induced (.ni) ^*RNAi*^SCoAS, Δ*idhm*/^*RNAi*^SCoAS and Δ*kdh-e2*/^*RNAi*^IDHm cell lines.

### α-Ketoglutarate is toxic if not metabolized at a high rate

As mentioned above, proline metabolism was strongly affected in the Δ*kdh-e2* mutant, with only production of α-ketoglutarate and 2-hydroxyglutarate (Fig 5A-B). As a consequence, growth of the Δ*kdh-e2* mutant was compromised in glucose-depleted medium containing 2 mM or 10 mM proline, while the addition of 10 mM malate or succinate rescued this growth phenotype (Fig 7A-B). However, addition of α-ketoglutarate at concentrations as low as 1 mM was detrimental for the survival of the Δ*kdh-e2* mutant (Fig 7A-B). This toxic effect of α-ketoglutarate was confirmed by the analysis of the ^*RNAi*^SDH.i and ^*RNAi*^SCoAS.i mutants, which are also affected in α-ketoglutarate metabolism. Indeed, growth of these two mutant cell lines was reduced in the presence of α-ketoglutarate (Fig 7B). It is also of note that expression of SCoAS was not fully abolished in the ^*RNAi*^SCoAS.i, which may explain the moderate reduction in its growth (Fig 7C). These data suggest that accumulation of α-ketoglutarate, or one of its metabolic derivatives, is toxic to PCF trypanosomes. Indeed, we cannot exclude that reduction of α-ketoglutarate to isocitrate, citrate or one of their metabolic products is responsible for this phenotype (see Fig 1D). To block the possible citrate/isocitrate production from α-ketoglutarate in the Δ*kdh-e2* and ^*RNAi*^SCoAS backgrounds, the *IDHm* gene was knocked-down or knocked-out to produce the Δ*kdh-e2*/^*RNAi*^IDHm and Δ*idhm*/^*RNAi*^SCoAS cell lines, respectively. Growth of these tetracycline-induced double mutants was strongly impaired by the addition of α-ketoglutarate as observed for the Δ*kdh-e2* and ^*RNAi*^SCoAS.i single mutants, while α-ketoglutarate stimulated growth of the Δ*idhm* as observed for the parental cell line (Fig 7A). This confirmed that it is accumulation of α-ketoglutarate itself that is responsible for this phenomenon.

It is noteworthy that accumulation of succinate was not toxic for trypanosomes, as exemplified by the absence of effect of 10 mM succinate on growth of the ^*RNAi*^SDH.i cell line (Fig 7B), which was no longer able to metabolize succinate (Fig 3B). As expected, malate stimulated growth of this mutant (Fig 7B).

### α-Ketoglutarate stimulates growth of epimastigote-like forms

To investigate whether TCA cycle intermediates are also carbon sources for the epimastigote trypanosomes, we took advantage of the *in vitro*-induced differentiation approach developed by Kolev *et al*., in which the differentiation of PCF to epimastigotes and then into metacyclics is triggered by overexpression of a single RNA binding protein (RBP6) [32]. In order to increase the proportion of epimastigotes, we selected a cell line expressing relatively low levels of RBP6, since the *in vitro*-induced differentiation of epimastigotes into metacyclics depends on strong overexpression of RBP6 [32]. Upon induction of RBP6 expression, the selected ^*OE*^RBP6.i cell line expressed the BARP (*brucei* alanine rich proteins) epimastigote differentiation marker [33], as well as the alternative oxidase (TAO), which has recently been described as strongly overexpressed in epimastigotes [34] (Fig 8A). This was confirmed by microscopy three days post induction, since 54% of the cells showed an epimastigote-like repositioning of the kinetoplast to reside close to the nucleus, while the other cells are procyclic-like (S4 Fig). In contrast no metacyclics were observed after eight days of induction, which is consistent by the stable expression of calflagin (Fig. 8A), a 10-fold upregulated flagellar protein in the bloodstream forms compared to procyclics [35]. We thus concluded that this tetracycline-induced ^*OE*^RBP6 cell line is enriched in epimastigote-like forms and/or forms in the process of becoming epimastigotes, whose carbon source requirements can be investigated. Growth of the ^*OE*^RBP6.i cells stopped after 6 days in the presence of 2 mM proline, while growth of non-induced ^*OE*^RBP6.ni cells was not affected. Interestingly, this growth defect is rescued by the addition of 10 mM proline or 10 mM α-ketoglutarate, but not the same quantity of succinate or malate. We confirmed that the expression profile of BARP and TAO is not affected by the presence of 10 mM α-ketoglutarate or 10 mM proline (Fig 8C) and ~50% of the cells showed an epimastigote-like repositioning of the kinetoplast 3 days post-induction. This data suggested that the amount of proline present in the midgut of the fly (1-2 mM) might be not sufficient to sustain growth of the epimastigotes and/or procyclic cells differentiating into epimastigotes and that additional carbon sources, such as α-ketoglutarate, should be needed to complete the development of the parasite *in vivo*.

**Fig. 8.**
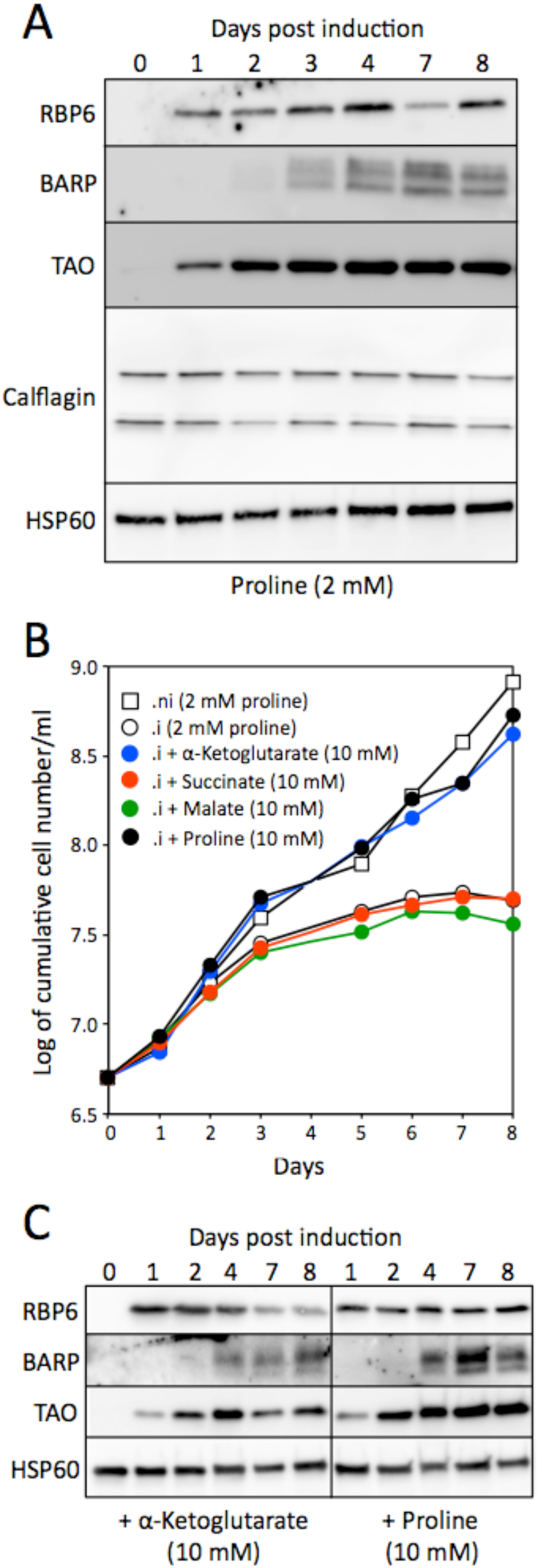
Large amounts of carbon sources are required for the growth of epimastigote-like cells. Panels A and C show Western blot analyses with the indicated immune sera (BARP, *brucei* alanine rich proteins; TAO, terminal alternative oxidase) of the ^*OE*^RBP6 cell line upon tetracycline induction and growth in low-proline conditions (2 mM) after addition (panel B) or not (panel C) of 10 mM α-ketoglutarate or 10 mM proline. HSP60 was used as loading control. Panel B shows growth curves of the tetracycline-induced (.i) and non-induced (.ni) ^*OE*^RBP6 cell line, in the presence of 2 mM proline, with or without addition of 10 mM α-ketoglutarate, succinate, malate, or proline. The curves are representative replicates of three different experiments.

## Discussion

Procyclic trypanosomes are thought to have developed a central metabolic network adapted to the metabolism of three main carbon sources for their energy, *i.e.* glucose, glycerol and proline [5,10,36]. Here we show that the parasite efficiently metabolizes a number of other carbon sources, including pyruvate, alanine and the TCA cycle intermediates succinate, malate and α-ketoglutarate. Several reports previously described the ability of African trypanosomes to consume and metabolize TCA cycle intermediates. For instance, α-ketoglutarate has been described to stimulate respiration and to sustain mobility of the stumpy forms of *T. brucei,* which are a growth-arrested transition form found in the bloodstream that are pre-adapted to differentiation into PCF [37–40]. In the 1960s, Riley showed that TCA cycle intermediates, including α-ketoglutarate, succinate, malate and fumarate, stimulate oxygen consumption of *T. brucei rhodesiense* culture forms (PCF) [36]. However, the metabolism of these TCA cycle intermediates by PCF trypanosomes has not been investigated so far in the context of the insect-like environment, that is to say in the absence of glucose but with proline present in the low millimolar range (1-2 mM), as described for the midgut of the tsetse fly [20]. Here we showed that in the presence of 2 mM proline, consumption and metabolism of succinate, malate and α-ketoglutarate takes place. More importantly, addition of 1-10 mM of any one of these three TCA cycle intermediates stimulates growth of the parasite and rescues growth of the parasite in the presence of 2 mM and 0.2 mM proline, respectively.

We took advantage of the unexpectedly high mitochondrial metabolic capacity developed by the PCF trypanosomes in the presence of α-ketoglutarate, succinate or malate to carry out a detailed analysis of the TCA cycle and its branched pathways. This allowed us to show (*i*) the high metabolic capacity of the malate/α-ketoglutarate branch of the TCA cycle, (*ii*) the toxicity of α-ketoglutarate intracellular accumulation and (*iii*) the production of 2-hydroxyglutarate from metabolism of α-ketoglutarate and proline. (*i*) Van Weelden *et al.* previously demonstrated that PCF trypanosomes cultured in rich medium do not need to oxidize glucose *via* a complete TCA cycle fed with glucose-derived acetyl-CoA [15]. Indeed, they showed that the growth rate of the Δ*aco* and parental cell lines was identical in the standard glucose-rich medium and, more importantly, the malate/α-ketoglutarate branch of the TCA cycle (steps 11-14 in Fig 1A) showed no significant activity in PCF trypanosomes, as measured by [^14^C]-CO_2_ release from labeled glucose. However, these data did not exclude a functional malate/α-ketoglutarate branch, which is used under specific nutritional conditions, or in particular developmental stages, and the sensitivity of the assay used in that work was limited. Here we have demonstrated the functionality of this metabolic branch by forcing the parasite to use it in the presence of 10 mM malate and 2 mM proline. In these conditions, malate is converted to succinate *via* both the reducing (steps 9-10 in Fig 1C) and oxidative (steps 11-14 and 5-6) branches of the TCA cycle. Indeed, fumarate production from malate is abolished in the ^*RNAi*^FHc/m.i mutant (step 10), while succinate production from malate is little affected, and succinate production is only reduced by 1.8-fold in the ^*RNAi*^FRDg/m.i mutant (step 9). These data can only be explained by a significant metabolic flux through the malate/α-ketoglutarate branch of the TCA cycle fed with extracellular malate. Since this branch of the TCA cycle does not seem to be important for their energy in standard culture conditions in the wild type PCF trypanosomes [15], its main function in the procyclic trypanosomes could be the production of citrate and/or isocitrate to supply other metabolic pathways. This hypothesis is consistent with the glycosomal localization of the IDHg isoform, which requires isocitrate for NADH and/or NADPH production within the organelle [41]. However, as opposed to most eukaryotes, trypanosomes do not use TCA-cycle derived citrate to produce the precursor of *de novo* fatty acid biosynthesis, *i.e.* acetyl-CoA, in the cytosol. The parasites lack the key enzyme of this pathway, *i.e.* cytosolic acetyl-CoA lyase, and instead use the cytosolic AMP-dependent acetyl-CoA synthetase to produce acetyl-CoA from acetate [42]. Interestingly, Dolezelova *et al*. recently took advantage of an *in vitro* differentiation assay based on RBP6 overexpression to show that all of the enzymes of the malate/α-ketoglutarate branch of the TCA cycle are strongly overexpressed upon differentiation into the epimastigote forms of *T. brucei* [34]. This suggests that this branch of the TCA cycle and/or the full TCA cycle is required for epimastigotes and/or during differentiation of procyclics into epimastigotes. (*ii*) Accumulation of the TCA cycle intermediate, α-ketoglutarate is toxic for PCF, while accumulation of succinate in the ^*RNAi*^SDH.i mutant cultivated in the presence of 10 mM succinate is not toxic. α-Ketoglutarate toxicity was deduced from the death of the Δ*kdh-e2* and Δ*kdh-e2*/^*RNAi*^IDHm.i mutants in the presence of α-ketoglutarate, which is not efficiently metabolized in these mutants compared to the parental cell line. α-Ketoglutarate is a key metabolite at the interface between metabolism of carbon and nitrogen [43], which has recently emerged as a master regulator metabolite in prokaryotes and cancer cells [44,45]. Consequently, intracellular accumulation of large amounts of α-ketoglutarate could affect several essential pathways, by mechanisms that are currently unknown. (*iii*) 2-Hydroxyglutarate is a five-carbon dicarboxylic acid occurring naturally in animals, plants, yeasts and bacteria. It has recently been described as an epigenetic modifier that governs T cell differentiation and plays a role in cancer initiation and progression [28,46]. This metabolite was also detected in the metabolome of BSF trypanosomes, being derived from the metabolism of glutamate [30]. Here we showed that 2-hydroxyglutarate represents 37% of the total end-products excreted by PCF from catabolism of α-ketoglutarate, with an excretion rate of 2-hydroxyglutarate 3.4 times higher than that of end-products from glucose metabolism (6.77 *versus* 2.02 μmole/h/mg of protein). This highlights the high capacity of the enzyme responsible for α-ketoglutarate reduction to 2-hydroxyglutarate. This reaction probably results from the promiscuous action of malate dehydrogenase on α-ketoglutarate or from oncogenic mutations in isocitrate dehydrogenase enzymes, as previously described in mammalian cells [29,47]. This NADH-consuming reaction compensates for the NADH-producing reactions involved in the production of other end-products excreted from α-ketoglutarate metabolism, and may therefore explain the very high α-ketoglutarate metabolic flux. Indeed, maintenance of the mitochondrial redox balance required for α-ketoglutarate metabolism is performed by fast-acting substrate level reactions, with little or no involvement of the respiratory chain, which operates at a lower rate compared to substrate level reactions. Most cells express 2-hydroxyglutarate dehydrogenase enzymes (2HGDH), which irreversibly catalyse the reverse oxidative reaction in order to prevent the loss of carbon moieties from the TCA cycle and would protect from the accumulation of 2-hydroxyglutarate [28]. The *T. brucei* genome contains a putatively annotated *2HGDH* gene (Tb927.10.9360), however since 2-hydroxyglutarate is produced in high quantities in PCF organisms its role in these cells is uncertain.

Trypanosomatids convert carbon sources to partially oxidized end-products that are excreted into the environment [5]. Some of these metabolites constitute good alternative carbon sources for the parasite, as exemplified by efficient metabolism of alanine by *T. cruzi*, while this amino acid is also excreted from glucose breakdown [48]. Here, we report that end-products excreted from the metabolism of glucose by PCF trypanosomes, such as succinate, alanine and pyruvate, are re-consumed after glucose has been used up. Indeed, ^13^C-enriched succinate and alanine excreted from catabolism of [U-^13^C]-glucose are re-consumed and converted to ^13^C-enriched acetate after glucose depletion. This phenomenon resembles the “acetate switch” which has been well described in prokaryotes, in which abundant or preferred nutrients, such as glucose, are first fermented to acetate, followed by the import and utilization of that excreted acetate to enhance survival of the cells [49]. This “acetate switch” occurs when cells deplete their environment of acetate-producing carbon sources.

We previously described that PCF trypanosomes cultivated in rich conditions use ~5 times more glucose than proline to feed their central carbon metabolism, and switch to proline metabolism in the absence of glucose by increasing its rate of consumption up to 5 fold [6,7]. Indeed, glucose is first fermented to excreted acetate, succinate, alanine and pyruvate, before switching to proline that is primarily metabolized in the mitochondrion with an increased contribution of the respiratory chain. Here we showed that this “proline switch” is accompanied by the re-utilization and conversion of glucose-derived end-products (succinate, pyruvate and alanine) to acetate. As opposed to bacteria or yeasts, however, acetate does not feed carbon metabolism of PCF and is the ultimate excreted end-product from the breakdown of the different carbon sources, including succinate and alanine. The ratio between the two main excreted end-products from glucose metabolism (acetate/succinate) has been reported to be between 0.3 and 4 in different studies [15,25], which has been interpreted to reflect a high flexibility of flux distribution between the acetate and succinate branches of the metabolic network [22,50]. In light of our observations, however, it appears that the conversion of excreted glucose-derived succinate to excreted acetate, following uptake and further metabolism of succinate, provides an alternative explanation for these heterogeneous data.

The *in vitro* differentiation model driven by overexpression of RBP6 was instrumental in showing that epimastigotes and/or cells in the process of differentiating into epimastigotes have a higher demand for carbon sources than procyclic trypanosomes to feed their central carbon metabolism. Indeed, a recent analysis of the proteome and metabolic capability showed that enzymes involved in proline catabolism, as well as mitochondrial respiratory capacity, are upregulated in the epimastigote-enriched population when compared to procyclic trypanosomes. This suggests an increased consumption of carbon sources, probably to meet an increased energy demand [34]. Consistent with these observations, our data demonstrated that growth of epimastigote-like cells, in contrast to that of procyclic cells, is affected by the presence of relatively small amounts of proline (2 mM), which corresponds to the level detected in the insect vector. Interestingly, adding 10 mM proline or 10 mM α-ketoglutarate restored growth, suggesting that this higher demand for carbon and energy sources may be supported by carbon sources other that proline, such as α-ketoglutarate. This increased demand for carbon/energy may contribute to the production of reactive oxygen species involved in the differentiation process of the parasite [34]. Alternatively, inhospitable organs of the fly or structures difficult to cross, such as the proventriculus, may require increased catabolic capacity. Indeed, the proventriculus is an active immune tissue of the insect that represents a hurdle to the spread of trypanosomes from the midgut to the salivary glands, since only a few trypanosomes can pass through it. Unfortunately, with the exception of amino acids [20], the content of metabolites in the tsetse midgut, including the proventriculus, has not been studied so far. TCA cycle intermediates could be present in significant amounts where, for example, tsetse resident symbionts such as *Sodalis glossinidius* metabolize N-acetylglucosamine and glutamine to produce partially oxidized end-products, which are released to the midgut lumen of the fly [18]. These may then promote trypanosome development. An exhaustive analysis of the metabolite content of the intestine of naive and infected insects is necessary to deepen our understanding of the role played by TCA cycle intermediates and other carbon sources in the development of trypanosomes in tsetse flies.

## Materials and Methods

### Trypanosomes and cell cultures

The procyclic form of *T. brucei* EATRO1125.T7T (TetR-HYG T7RNAPOL-NEO) and AnTat 1.1 were cultured at 27°C in SDM79 medium containing 10% (v/v) heat inactivated fetal calf serum and 5 μg/ml hemin [51] and in the presence of hygromycin (25 μg/ml) and neomycin (10 μg/ml). All mutant cell lines have initially been produced and cultivated in the standard SDM79 medium. Alternatively, the cells were cultivated in a glucose-depleted medium derived from SDM79 supplemented with 50 mM N-acetylglucosamine, a specific inhibitor of glucose transport that prevents consumption of residual glucose [13]. Control growth conditions in the presence of glucose were performed in glucose-depleted conditions, in which the glucose transport inhibitor N-acetylglucosamine was omitted and 10 mM glucose was added. The growth was followed by counting the cells daily with a Guava EasyCyte™ cytometer. The Alamar Blue assay was used to study the effect of metabolites on parasite growth. To do this, cells at a final density of 2 × 10^6^ cells/ml were diluted in 200 μl of glucose-depleted medium supplemented with 6 mM proline/10 mM glucose, 2 mM proline or 12 mM proline containing 1 μM to 100 mM of the analyzed metabolite and incubated for 48 h at 27°C in microplates, before adding 20 μl of 0.49 mM Alamar Blue (Resazurin). Measurement of fluorescence was performed with the microplate reader Fluostar Optima (BMG Labtech) at 550 nm excitation wavelength and 600 nm emission wavelength as previously described [23].

### Inhibition of gene expression by RNAi

The inhibition by RNAi of gene expression in the PCF trypanosomes was performed by expression of stem-loop “sense/antisense” (SAS) RNA molecules of the targeted sequences introduced into the pLew100 or a single fragment in the p2T7^Ti^-177 expression vectors (kindly provided by E. Wirtz and G. Cross and by B. Wickstead and K. Gull, respectively) [52,53]. Plasmids pLew-FRDg/m1-SAS, p2T7-FHc/m-SAS, pLew-SDH-SAS, pLew-PDH-E2-SAS, pLew-AAT-SAS, p2T7-PRODH were used to generate the ^*RNAi*^FRDg/m1-B5 [24], ^*RNAi*^FHc/m-F10 [25], ^*RNAi*^SDH [7], ^*RNAi*^PDH-E2 [7], ^*RNAi*^AAT [11] and ^*RNAi*^PRODH [6] cell lines, as previously reported [54]. The ^*RNAi*^AAT and ^*RNAi*^PRODH mutants produced in the 29-13 cell line (derived from strain 427) and the other RNAi cell lines obtained in the EATRO1125.T7T background were grown in SDM79 medium containing hygromycin (25 μg/ml), neomycin (10 μg/ml) and phleomycin (5 μg/ml). A 591 bp fragment of the beta subunit of SCoAS (Tb927.10.7410) was PCR amplified and cloned into the p2T7^Ti^-177 plasmid using the BamHI and HindIII restriction sites (p2T7^Ti^-177-SCoAS). Procyclic EATRO1125.T7T cells were transfected with the p2t7-177-SCoAS plasmid to generate the ^*RNAi*^SCoAS cell line. To assemble p2T7^Ti^-177_IDHm^RNAi^(ble), a 1127 bp fragment of IDHm (Tb927.8.3690) was amplified (primers IDHm_fwd: TGTCTACAACACGTCCAA and IDHm_rev BamHI: CGATAggatccGATGGTTTTGATCGTTGC) from genomic AnTat1.1 DNA and cloned into the p2T7^Ti^-177 plasmid using the BamHI and HindIII restriction sites.

### Production of null mutants

The *ASCT* gene was replaced by the blasticidin/puromycin resistance markers to generate the previously reported Δ*asct* null mutant [14]. The Δ*ach*/^*RNAi*^ASCT cell line was generated by introducing the pLew-ASCT-SAS plasmid in the Δ*ach* background [14]. To delete the genes encoding the subunit E2 of KDH (KDH-E2), the resistance markers blasticidin (BLA) and puromycin (PAC) were PCR amplified using long primers with 80 bp corresponding to the 5’UTR and 3’UTR region of the *KDH-E2* gene (Tb927.11.9980). To replace the two *KDH-E2* alleles, the EATRO1125.T7T PCF was transfected with 10 μg of purified PCR products encoding the resistance markers flanked by UTR regions. The selected Δ*kdh-e2*::*PAC/*Δ*kdh-e2*::*BLA* cell line was named Δ*kdh-e2*. The cell line Δ*kdh-e2*/^*RNAi*^IDHm was generated by transfection of p2T7^Ti^-177_IDHm^RNAi^(ble) into Δ*kdh-e2* and selection with phleomycin. For generation of a homozygous Δ*aco* cell line in the EATRO1125.T7T background, previously reported targeting constructs [15] were modified by replacing the neomycin and hygromycin selection markers with blasticidin and puromycin, respectively. For homozygous deletion of *IDHm* (Tb927.8.3690), the EATRO1125.T7T line was transfected with targeting constructs having the blasticidin and puromycin selection marker cassettes flanked by *IDHm* 5’-UTR (797 bp) and 3’-UTR (876 bp) sequences, amplified from genomic DNA of strain MiTat1.4. The selected Δ*idhm*::*PAC/*Δ*idhm*::*BLA* cell line was named Δ*idhm*. The Δ*idhm* null mutant was transfected with the p2T7^Ti^-177-SCoAS plasmid to generate the Δ*idhm*/^*RNAi*^SCoAS cell line.

### *In vitro* differentiation of PCF expressing RBP6

The *RBP6* gene was amplified by PCR and cloned via HindIII/BamHI into the pLew100v5b1d vector (pLew100v5 modified with a blasticidin resistance gene *BSD*) [23]. The EATRO1125.T7T cell line was transfected with the pLew100v5-RBP6 linearized with NotI in pools to generate the ^*OE*^RBP6. *In vitro* differentiation experiments were done as described in [23,32] in SDM79 medium without glucose in the presence of 50 mM *N*-acetyl-D-glucosamine and 10% (v/v) heat-inactivated fetal calf serum.

### Analysis of excreted end-products from the metabolism of carbon sources by proton NMR

2 × 10^7^ *T. brucei* PCF cells were collected by centrifugation at 1,400 g for 10 min, washed once with phosphate-buffered saline (PBS) and incubated in 1 ml (single point analysis) of PBS supplemented with 2 g/l NaHCO_3_ (pH 7.4). For kinetic analysis, 1 × 10^9^ cells were incubated in 15 ml under the same conditions. Cells were maintained for 6 h at 27°C in incubation buffer containing one or two ^13^C-enriched or non-enriched carbon sources. The integrity of the cells during the incubation was checked by microscopic observation. The supernatant (1 ml) was collected and 50 μl of maleate solution in D_2_O (10 mM) was added as internal reference. H-NMR spectra were performed at 500.19 MHz on a Bruker Avance III 500 HD spectrometer equipped with a 5 mm cryoprobe Prodigy. Measurements were recorded at 25°. Acquisition conditions were as follows: 90° flip angle, 5,000 Hz spectral width, 32 K memory size, and 9.3 sec total recycle time. Measurements were performed with 64 scans for a total time close to 10 min 30 sec. Resonances of the obtained spectra were integrated and metabolites concentrations were calculated using the ERETIC2 NMR quantification Bruker program. The identification of 2-hydroxyglutarate in some samples was duly confirmed from H-NMR analyses carried out at 800 MHz, after spiking with the pure compound.

### Western blot analyses

Total protein extracts (5 × 10^6^ cells) were separated by SDS-PAGE (10%) and immunoblotted on TransBlot Turbo Midi-size PVFD Membranes (Bio-Rad) [55]. Immunodetection was performed as described [55,56] using as primary antibodies, the rabbit anti-ASCT (1:500) [57], anti-aldolase (1:10,000) [58], anti-PPDK (1:500) [54], anti-ENO (1:100,000, gift from P. Michels, Edinburgh, UK), anti-FRD (1:100) [24], anti HSP60 (1:10,000) [59], anti-RBP6 (1:500, gift from C. Tschudi, New Haven, USA), anti BARP (1:2,500, gift from I. Roditi, Bern, Switzerland) [33], anti-calflagin (1:1,500) or the mouse anti-PDH-E2 (1:500) [21], anti-TAO 7D3 (1:100, gift from M. Chaudhuri, Nashville, USA) [60] and anti-FH (1:100) [25]. Rabbit anti-IDHm was raised against recombinant His-tagged full length *T. brucei* IDHm expressed in and purified from *E. coli*. After intradermal immunization with Freund’s complete adjuvant, 12 boosts were required until final bleeding after 180 days (custom immunization by Pineda, Berlin). The recombinant full length SCoAS subunit β with N-terminal 6x His tag was affinity-purified from *E. coli* under native conditions and sent to Davids Biotechnology (Regensburg, Germany) for polyclonal antibody production. Anti-rabbit or anti-mouse IgG conjugated to horseradish peroxidase (Bio-Rad, 1:5,000 dilution) were used as secondary antibody and detected using the Clarity Western ECL Substrate as described by the manufacturer (Bio-Rad). Images were acquired and analyzed with the ImageQuant LAS 4000 luminescent image analyzer.

### Enzymatic activity assays

For PRODH and SDH activities, Log phase PCF cells were harvested and washed twice with STE buffer (25 mM Tris-HCl, pH 7.4, 1 mM EDTA, 0.25 M sucrose, Protease inhibitors) and treated with 0.35 mg digitonin per mg of protein during 4 min at room temperature. After centrifugation 2 min at 12,000 x g, the enzymatic activities were determined in the pellets resuspended in STE. Proline dehydrogenase (PRODH) and succinate dehydrogenase activities (SDH) were measured following the reduction of the electron-accepting dye dichlorophenolindophenol (DCPIP) at 600 nm [61]. The assays contained 11 mM MOPS pH 7.5, 11 mM MgCl_2_, 11% glycerol, 56 μM DCPIP, 0.9 mM PMS, 10 mM of proline for PRODH activity or 20 mM of succinate for SDH activity. The alanine aminotransferase (AAT) activity was measured following the oxidation of NADH at 340 nm [62]. The malate dehydrogenase (MDH) activity was determined as a control [63].

## Acknowledgements

We thank Paul A. Michels (Edinburgh, Scotland), Christian Tschudi (New Haven, USA), Minu Chaudhuri (Nashville, USA) and Isabel Roditi (Bern, Switzerland) for providing us with the anti-enolase, anti-RBP6, anti-TAO and anti-BARP immune sera, respectively. F. Bringaud’s group was supported by the Centre National de la Recherche Scientifique (CNRS), the Université de Bordeaux, the ANR through the grants GLYCONOV (grant number ANR-15-CE15-0025-01) and ADIPOTRYP (grant number ANR-19) and the Laboratoire d’Excellence (LabEx) ParaFrap ANR-11-LABX-0024. M. Barrett’s group was supported by the Wellcome Trust core grant to the Wellcome Centre of Integrative Parasitology (grant number 104111/Z/14/Z). A. M. Silber’s group is supported by FAPESP (grant number 2016/06034-2) and CNPq (grants number 308351/2013-4 and 404769/2018-7). A. M. Silber and M. Barrett are funded by a joint FAPESP-MRC/UKRI-NEWTON FUND award: “Bridging epigenetics, metabolism and cell cycle in pathogenic trypanosomatids” (grant number 2018/14432-3), A. Zikova’s group by Grant Agency of the Czech Republic (20-14409S) and ERD fund (CZ.02.1.01/0.0/0.0/16_019/0000759) and M. Boshart’s group was funded by DFG SPP1131, and BayFrance agency supported collaboration between the Bordeaux and Munich labs. MetaboHub-MetaToul (Metabolomics & Fluxomics facilities, Toulouse, France, http://www.metatoul.fr) is supported by the ANR grant MetaboHUB-ANR-11-INBS-0010. JCP is grateful to INSERM for funding a temporary full-time researcher position.

---

**S1 Fig.**
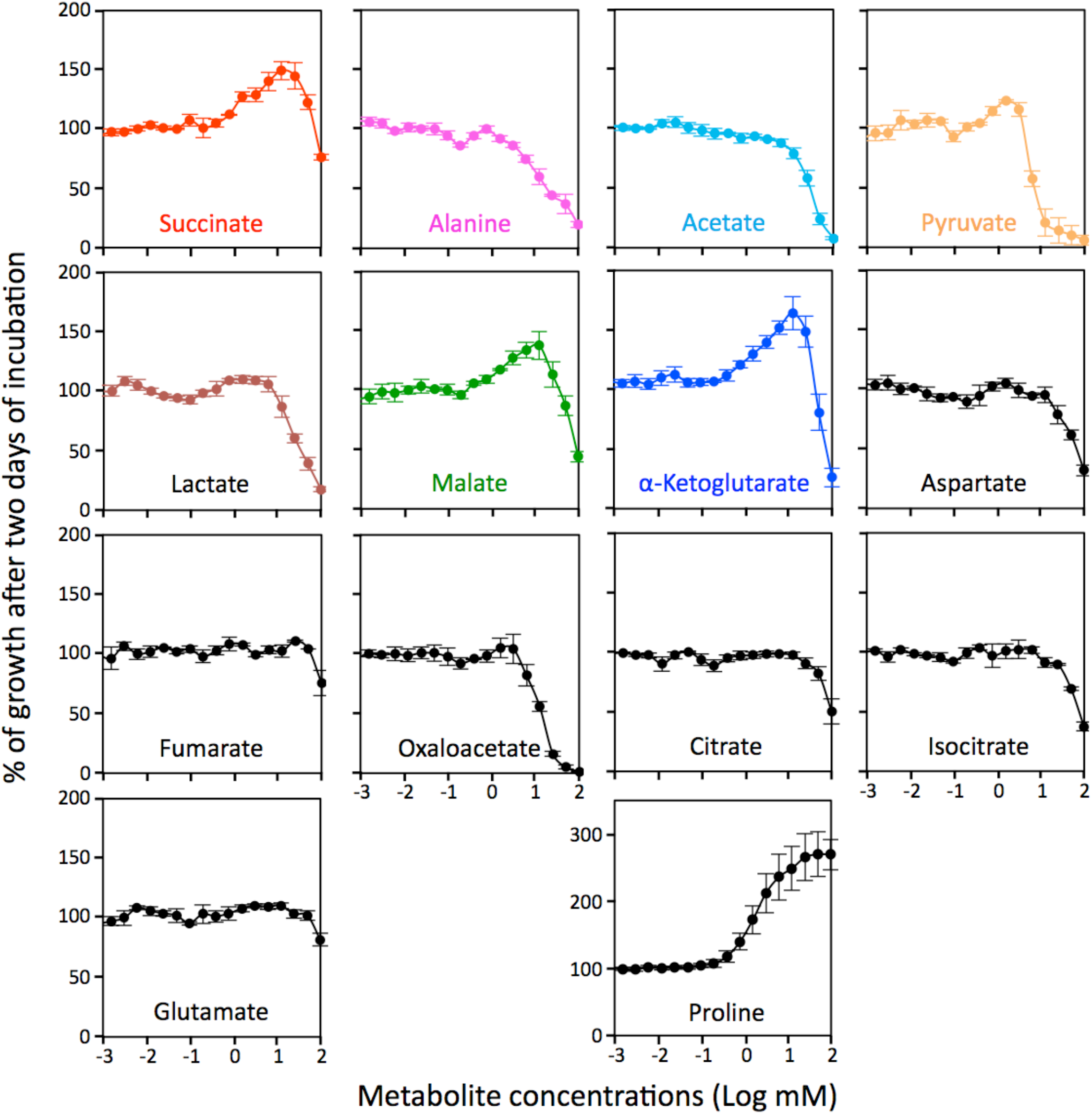
Growth of the PCF trypanosomes in the glucose-depleted SDM79 medium containing 2 mM proline in the presence of added 10 μM to 100 mM metabolite, using the Alamar Blue assay. Incubation was started at 2 x 10^6^ cell density and the Alamar Blue assay was performed after 48 h at 27°C as described before [23].

**S2 Fig.**
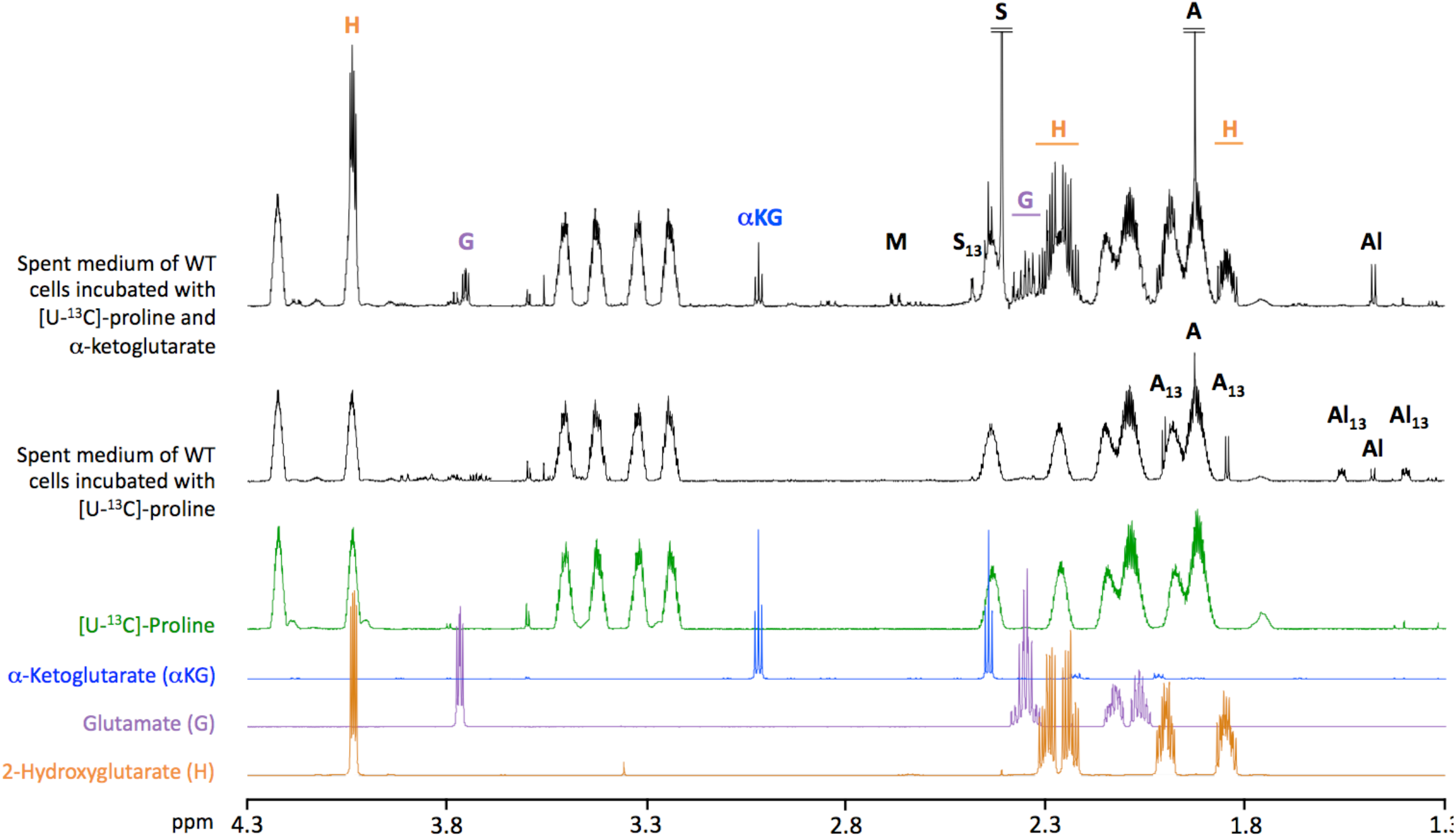
H-NMR analysis of samples (black) and controls (colored) performed at 800 MHz to identify acetate (A), alanine (Al), glutamate (G), 2-hydroxyglutarate (H), α-ketoglutarate (αKG), malate (M), proline and succinate (S). The resonances corresponding to ^13^C-enriched molecules are indicated in index.

**S3 Fig.**
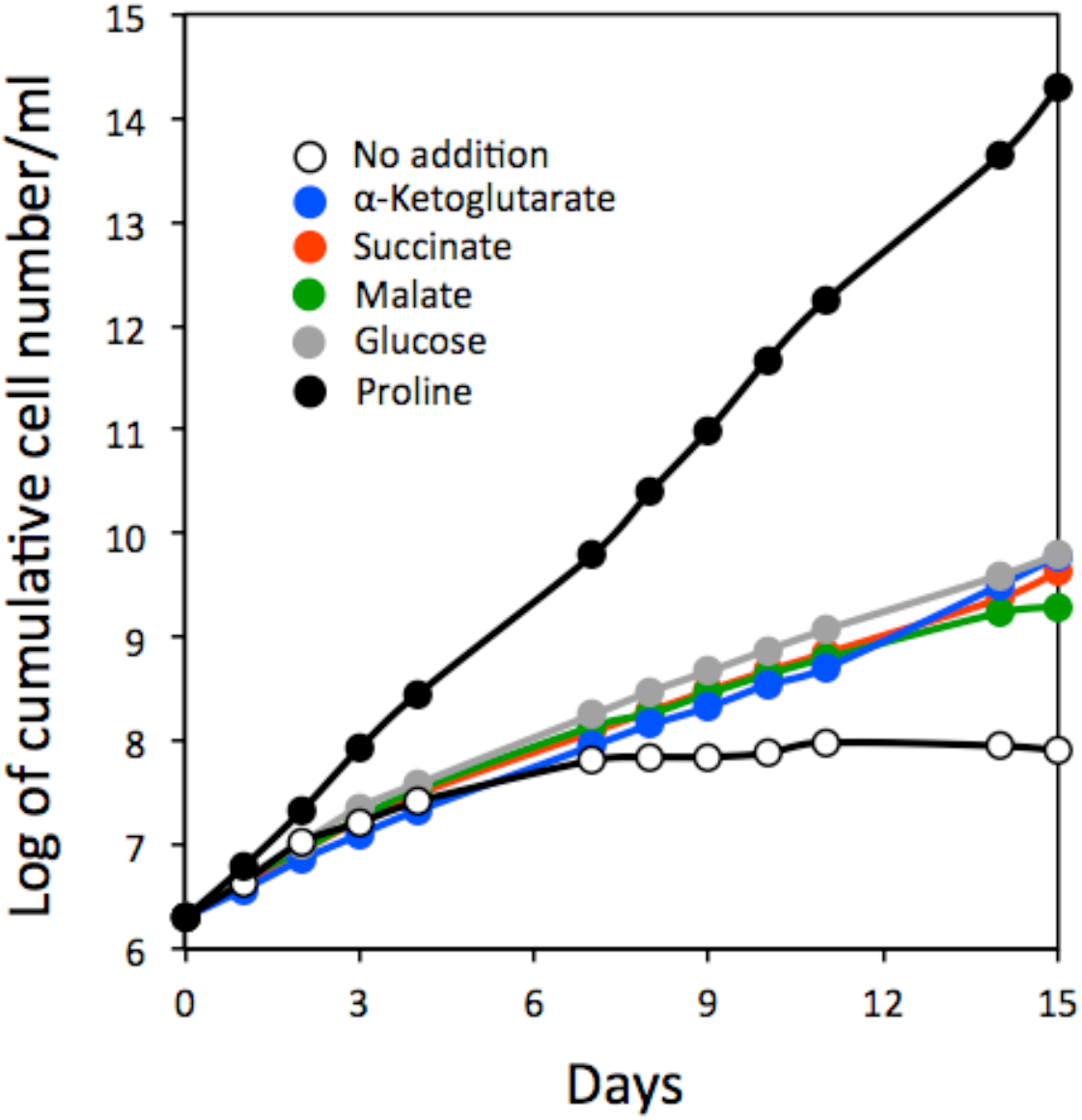
Growth curves of PCF trypanosomes grown in low-proline conditions (0.2 mM) in the presence or absence of 10 mM proline, glucose, α-ketoglutarate, succinate or malate. Cells were maintained in the exponential growth phase (between 10^6^ and 10^7^ cells/ml), and cumulative cell numbers reflect normalization for dilution during cultivation.

**S4 Fig.**
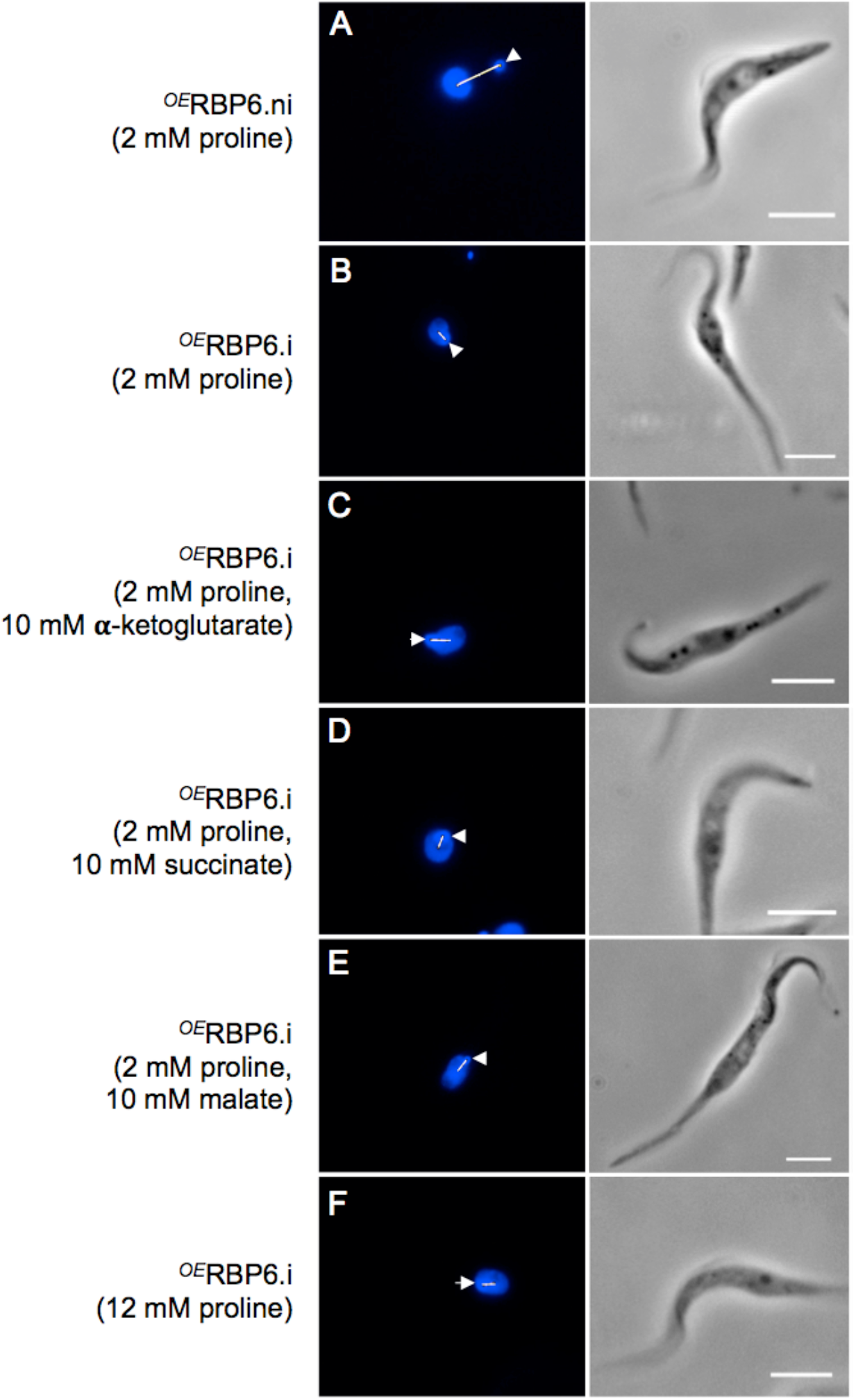
Imaging of procyclic and epimastigote-like forms. Illustration of microscopic analyses of non-induced (.ni) (A) or induced (.i) (B-F) ^*OE*^RBP6 cells grown in the presence of 2 mM proline complemented or not (A-B) with 10 mM of α-ketoglutarate (C), succinate (D), malate (E) or proline (F). The non-induced population is composed of procyclic trypanosomes (A), while epimastigote-like cells mainly composed the induced population regardless of the culture conditions. DAPI staining of DNA is shown on the left panels, in which kinetoplasts are highlighted by arrowheads and the distance between kinetoplasts and nuclei are shown by white lines, while the right panels show phase contrast (calibration bar: 5 μm). These analyses were performed three days post induction (B-F).

**S1 Table.**
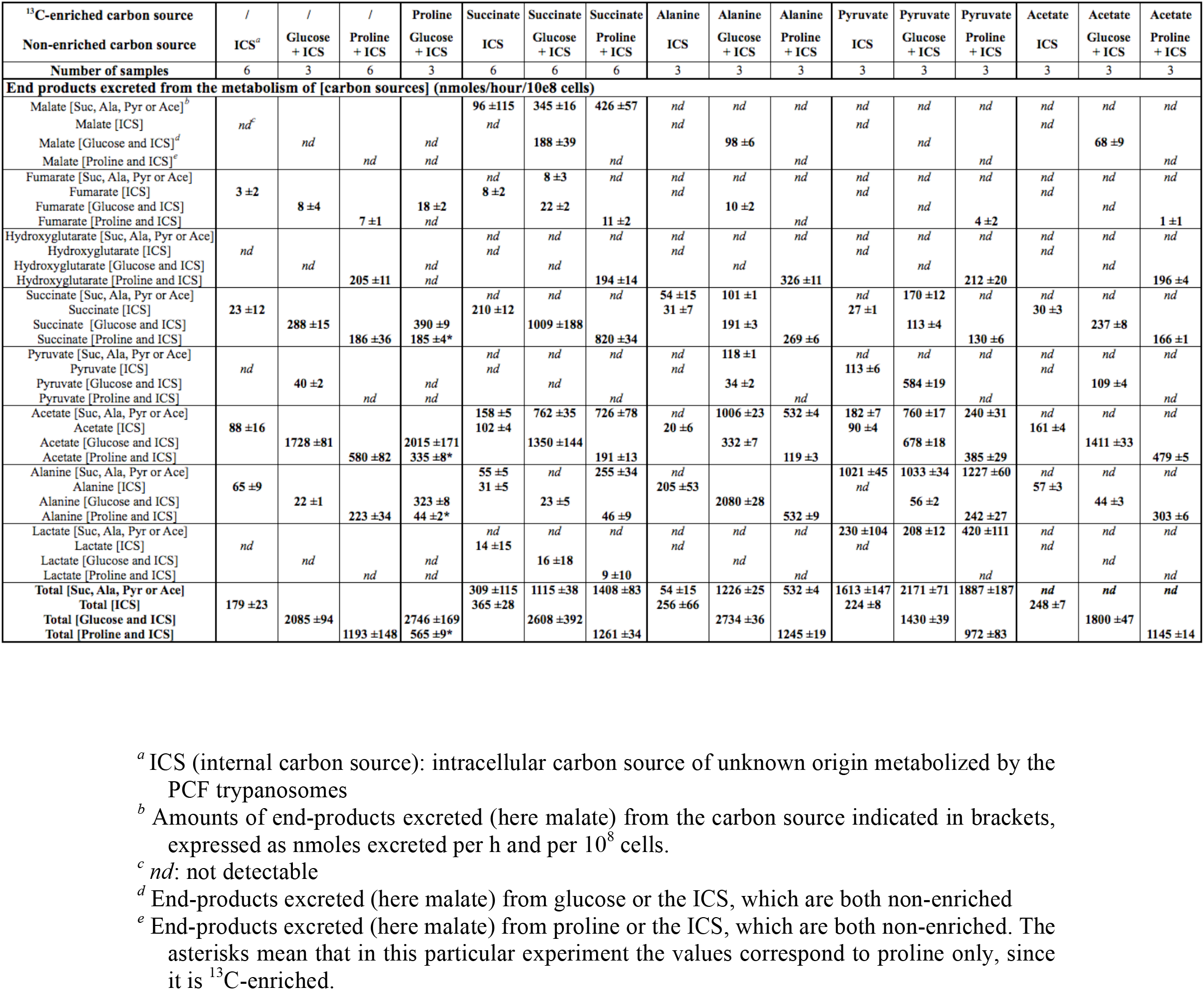
Excreted end-products from the metabolism of carbon sources in the PCF trypanosomes. The parasites were incubated with 4 mM [U-^13^C]-succinate, [U-^13^C]-alanine, [U-^13^C]-pyruvate or [U-^13^C]-acetate in the presence or absence of 4 mM glucose or proline.

